# Formation of polarized contractile interfaces by self-organized Toll-8/Cirl GPCR asymmetry

**DOI:** 10.1101/2020.03.16.993758

**Authors:** Jules Lavalou, Qiyan Mao, Stefan Harmansa, Stephen Kerridge, Annemarie C. Lellouch, Jean-Marc Philippe, Stephane Audebert, Luc Camoin, Thomas Lecuit

## Abstract

During development, interfaces between cells with distinct genetic identities elicit signals to organize local cell behaviors driving tissue morphogenesis. The *Drosophila* embryonic axis extension requires planar polarized enrichment of Myosin-II powering oriented cell intercalations. Myosin-II levels are quantitatively controlled by G protein-coupled receptor (GPCR) signaling whereas Myosin-II polarity requires patterned expression of several Toll receptors. How Toll receptors polarizes Myosin-II, and how this involves GPCRs, remain unknown. Here we report that differential expression of a single Toll receptor, Toll-8, polarizes Myosin-II via a novel binding partner, the adhesion GPCR Cirl/Latrophilin. Asymmetric expression of Cirl is sufficient to enrich Myosin-II and Cirl localization is asymmetric at Toll-8 expression boundaries. Exploring the process dynamically, we reveal that Toll-8 and Cirl exhibit mutually dependent planar polarity in response to quantitative differences in Toll-8 expression between neighboring cells. Collectively, we propose that a novel cell surface protein complex Toll-8/Cirl self-organizes to generate local asymmetric interfaces essential for planar polarization of contractile interfaces.

## Introduction

Dynamic cell behaviors that drive tissue morphogenesis are often organized at local cell interfaces. At one end, surface signaling between cells with distinct genetic identities via compartmentalized expression of ligands and receptors can generate mechanically activated interfaces in developmental (Dahmann et al., 2011) or pathological contexts (Bielmeier et al., 2016). For instance, spatial domains of Eph-Ephrin or Leucine-rich Repeats (LRRs) proteins induce mechanical barriers that prevent cell mixing in vertebrate and invertebrate models (Dahmann et al., 2011; Fagotto et al., 2013; Karaulanov et al., 2006; Milán et al., 2001; Paré et al., 2019; Smith and Tickle, 2006; Tomás et al., 2011). At the other end, surface signaling can also orient mechanical interfaces in the context of tissue planar polarity. Surface proteins in the core Planar Cell Polarity (PCP) pathway, such as Flamingo/Celsr and Frizzled, translate tissue-scale cues into vectorial cell polarity known to control planar polarized actomyosin contractility (Aw and Devenport, 2017; Nishimura et al., 2012). Tissue-level gradients of the Fat/Dachsous adhesion molecules in the Fat-PCP pathway induce planar polarized accumulation of the myosin Dachs at cell interfaces to drive dynamic cell rearrangements (Bosveld et al., 2012).

The early *Drosophila* embryo is an excellent system to investigate how cell groups with distinct genetic identities generate planar polarized mechanical interfaces. In the ventrolateral ectoderm, Myosin-II (Myo-II) is enriched at vertical interfaces between anteroposterior (AP) neighbors, which produces polarized actomyosin contractility powering AP axis extension (Bertet et al., 2004; Blankenship et al., 2006; Irvine and Wieschaus, 1994; Zallen and Wieschaus, 2004). The amplitude and polarity of actomyosin contractility appears to be controlled by diverse cell surface proteins. Levels of Myo-II activation at cell interfaces are quantitatively controlled by G protein-coupled receptor (GPCR) signaling (Garcia De Las Bayonas et al., 2019; Kerridge et al., 2016), whereas the polarized enrichment of Myo-II between AP neighbors appears to be governed by several Toll receptors, Toll-2,6,8 (Paré et al., 2014). Interactions between pair-rule genes define periodic and partially overlapping stripes of Toll receptors perpendicular to the AP axis (Paré et al., 2014). Thus, each column of cells expresses a different combination of Toll-2,6,8, which are thought to be collectively required for planar polarized Myo-II activity (Paré et al., 2014; Tetley et al., 2016). Toll receptors belong to the LRR super family and are well known for their conserved function in developmental patterning and innate immunity (Anthoney et al., 2018). They recently emerged as conserved molecules involved in embryonic axis elongation (Benton et al., 2016; Paré et al., 2014) and organogenesis (Yagi et al., 2010). How Toll receptors define polarized mechanical interfaces, and whether they interact with other cell surface receptors such as GPCRs, remain largely unexplored.

Here we investigated how Toll receptors control Myo-II planar polarity in conjugation with GPCR signaling in the *Drosophila* embryonic ectoderm and the larval wing disc epithelium. In contrast to the combinatorial Toll code model, we show that a single Toll receptor, Toll-8, enriches Myo-II at its expression boundary. This enrichment does not require the cytoplasmic tail of Toll-8. Further, we identified a novel Toll-8 binding partner, the adhesion GPCR Cirl/Latrophilin, which is required for Toll-8-mediated Myo-II enrichment both in embryos and wing discs. We show in wing discs that Cirl asymmetry is sufficient to enrich Myosin-II and we observed an interfacial asymmetry in Cirl apicobasal localization at Toll-8 expression boundaries. Moreover, when neighboring cells express different levels of Toll-8 in wing discs, both Toll-8 and Cirl exhibit robust and mutually dependent planar polarity. Our study thus reveals that Toll-8 and Cirl form a cell surface protein complex essential for planar polarized actomyosin activity, generate local asymmetric interfaces and co-polarize in a self-organized manner.

## Results

### Asymmetric expression of a single Toll receptor leads to Myo-II polarization in embryos

In the *Drosophila* embryonic ectoderm, it has been hypothesized that *trans*-interactions between different Toll receptors across cell-cell interfaces signal to polarize junctional Myo-II (Paré et al., 2014). In light of the observation that ectopic expression of Toll-2 or Toll-8 alone induces Myo-II enrichment late in embryogenesis (Paré et al., 2014), we first tested the simplest hypothesis that a single Toll is sufficient to polarize junctional Myo-II in the embryonic ectoderm. To this end, we injected embryos with dsRNAs targeting Toll-2,6,7 (*toll-2,6,7* RNAi), leaving only endogenous Toll-8 expressed in vertical stripes (Figure S1A), and observed Myo-II with mCherry-tagged Myo-II regulatory light chain (MRLC–Ch). We found Myo-II specifically enriched at the interfaces between Toll-8 expressing and non-expressing cells (Figures S1A and S1B), suggesting that asymmetric expression of a single Toll, Toll-8, leads to Myo-II enrichment independent of other Toll receptors.

To assess the sufficiency of Toll-8 asymmetry in Myo-II polarization independent of the anteroposterior (AP) patterning system, we engineered embryos expressing a single stripe of Toll-8 along the AP axis that runs orthogonal to the endogenous Toll-2,6,8 stripes, using an *intermediate neuroblasts defective* (*ind*) enhancer (Stathopoulos and Levine, 2005) (*ind-Toll-8–HA*, Figure 1A). We monitored Myo-II with GFP-tagged Myo-II regulatory light chain (MRLC–GFP) and detected a Myo-II enriched cable at the ventral boundary between ectopic Toll-8 expressing and wild-type cells (Figures 1A and 1B, Toll-8^FL^). Note that this ectopic Myo-II cable runs along the AP axis and is thus perpendicular to the endogenous planar polarity of Myo-II. By contrast, no Myo-II enrichment is detected at homotypic Toll-8 interfaces within the ind-Toll-8 domain (Figure 1C). When we deleted the Leucine-rich Repeats (LRRs) from the Toll-8 extracellular domain, Toll-8 was no longer localized to the plasma membrane and failed to accumulate Myo-II (Figures 1A, 1B and 1D, Toll-8^ΔLRR^), suggesting that the LRRs of Toll-8 are essential for Myo-II polarization. Consistent with an upregulation of cortical tension, the boundary of Toll-8 expressing cells was smoother when full-length Toll-8 (Toll-8^FL^) was expressed compared to Toll-8^ΔLRR^ (Figures 1B and 1E; smoothness defined by the ratio of distance between terminal vertices over total junctional length).

**Figure 1.**
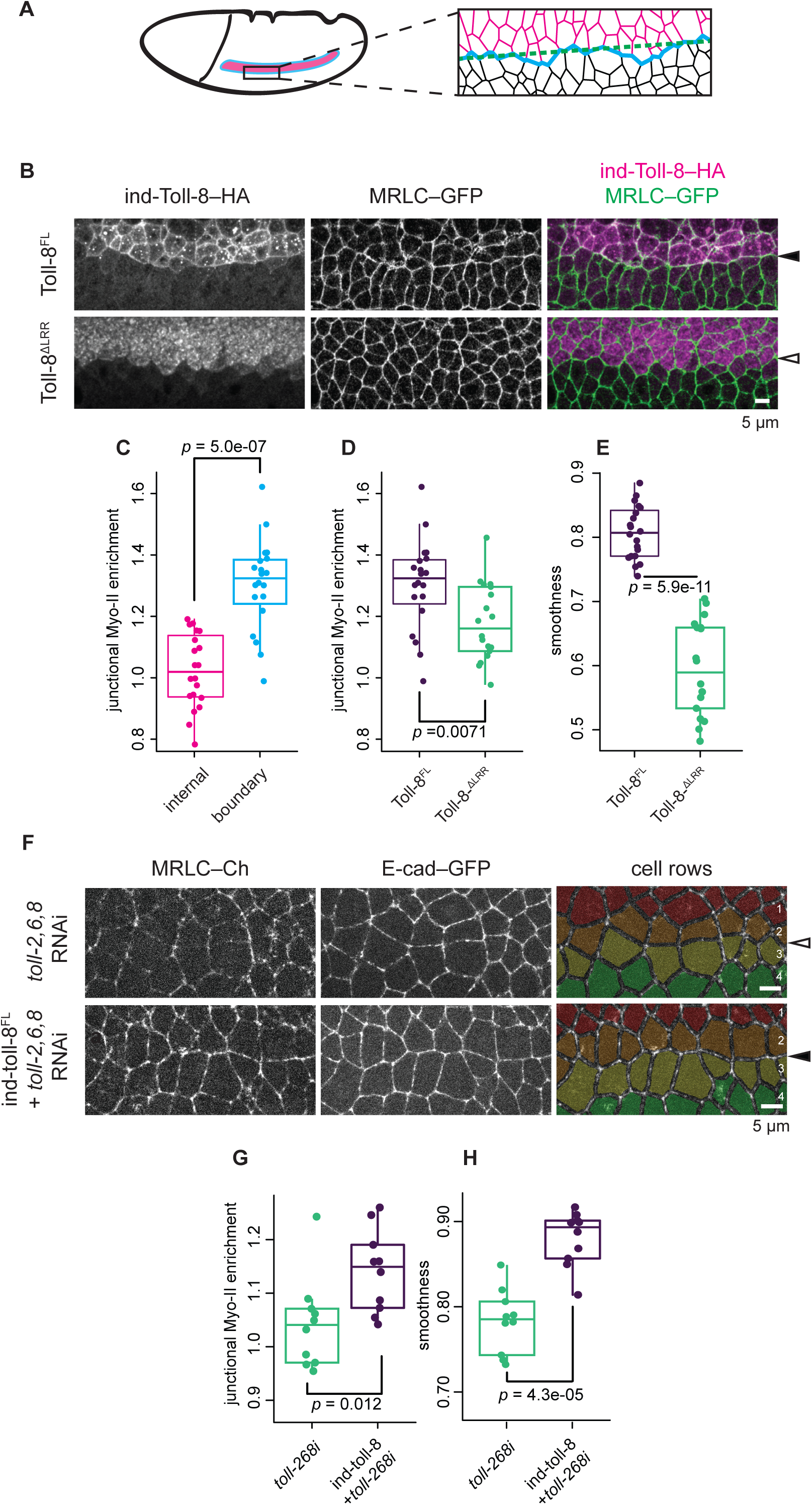
Toll-8 asymmetry leads to Myo-II enrichment in embryos, independent of other Toll receptors. **(A)** Schema showing an embryo expressing ind-Toll-8–HA (left, pink stripe) and a detailed view of cell interfaces around the ventral boundary of the ind-Toll-8–HA stripe (right). Interfaces within the ind-Toll-8–HA stripe in pink; those within the wild-type tissue in black; those at the ventral boundary of the ind-Toll-8–HA stripe in blue. Dashed line in green denotes the distance between the first and last vertices of the ind-Toll-8–HA ventral boundary. **(B)** Anti-Toll-8–HA (left) and anti-MRLC-GFP (middle) signals in *Drosophila* stage 7 embryos expressing full length Toll-8 (Toll-8^FL^, top, *n*=20) or Toll-8 with the extracellular LRRs removed (Toll-8^ΔLRR^, bottom, *n*=18) under the control of the *ind* promoter. Myo-II is enriched at the ventral boundary of the ind-Toll-8^FL^–HA expression domain (filled arrowhead) but not at the ind-Toll-8^ΔLRR^–HA boundary (empty arrowhead). **(C)** Junctional Myo-II enrichment relative to wild-type tissue within the ind-Toll-8^FL^–HA stripe (pink) or at the ventral boundary of the ind-Toll-8^FL^–HA stripe (blue) for the condition shown in (**B**, top). **(D and E)** Junctional Myo-II enrichment (**D**) and boundary smoothness (**E**) at the ventral *ind* boundary for the conditions shown in (**B**). **(F)** Still images from time lapse movies in *wt* embryos (top) or embryos expressing ind-Toll-8^FL^–HA (bottom) injected with dsRNAs against Toll-2,6,8 (*n*=10 each). Pseudo colors mark 4 horizontal cell rows. Myo-II is still enriched at the interface between cell rows 2 and 3 corresponding to the *ind* ventral boundary in the absence of endogenous Toll-2,6,8 (filled arrowhead). **(G and H)** Junctional Myo-II enrichment (**G**) and boundary smoothness (**H**) at the ventral *ind* boundary for the conditions shown in (**F**). Scale bars: 5 µm. Statistics: Mann-Whitney U test.

When we injected *ind-Toll-8–HA* embryos with dsRNAs targeting only endogenous Toll-2,6,8 (*toll-2,6,8* RNAi), Myo-II was still enriched at the boundary of ectopic Toll-8 expressing cells (Figures 1F, 1G, S1C and S1D). This boundary was also smooth (Figures 1F, 1H, S1C and S1D). Altogether, we conclude that an interface defined by asymmetric expression of a single Toll, Toll-8, is sufficient to polarize Myo-II.

### Asymmetric expression of a single Toll receptor polarizes Myo-II in wing discs

We wished to further dissect the capacity of Toll-8 to elicit junctional Myo-II enrichment using clonal analysis to generate random interfaces between wild-type and Toll-8 overexpressing cells. We chose to study Toll-8 in larval wing imaginal discs, since clonal analysis is not possible in early embryos. Wing imaginal discs exhibit polarized supracellular cables of Myo-II at the periphery of the pouch region near the hinge (LeGoff et al., 2013). Toll receptor overexpression causes wing developmental defects (Yagi et al., 2010) and Toll-8 is expressed in the wing hinge region but not in most of the pouch region (Alpar et al., 2018; Yagi et al., 2010). We induced clones overexpressing Toll-8–GFP in the larval wing disc epithelium and monitored Myo-II localization with MRLC–Ch 24 hours after clone induction in the wing pouch (Figure 2A). This resulted in striking junctional Myo-II enrichment specifically at the boundary of Toll-8 overexpressing clones, as no Myo-II enrichment is detected at homotypic Toll-8 interfaces within the Toll-8 clone (Figures 2A-2C). As in embryos, we found that LRRs were required for Toll-8 plasma membrane localization and Myo-II polarization in wing discs (Figures 2D and 2E, Toll-8^ΔLRR^). Toll-8^FL^ clones were more compact with a smoother boundary compared with Toll-8^ΔLRR^ clones (Figures 2D and 2F; smoothness defined by the mean angle between neighboring vertices as shown in inset of Figure 2A).

**Figure 2.**
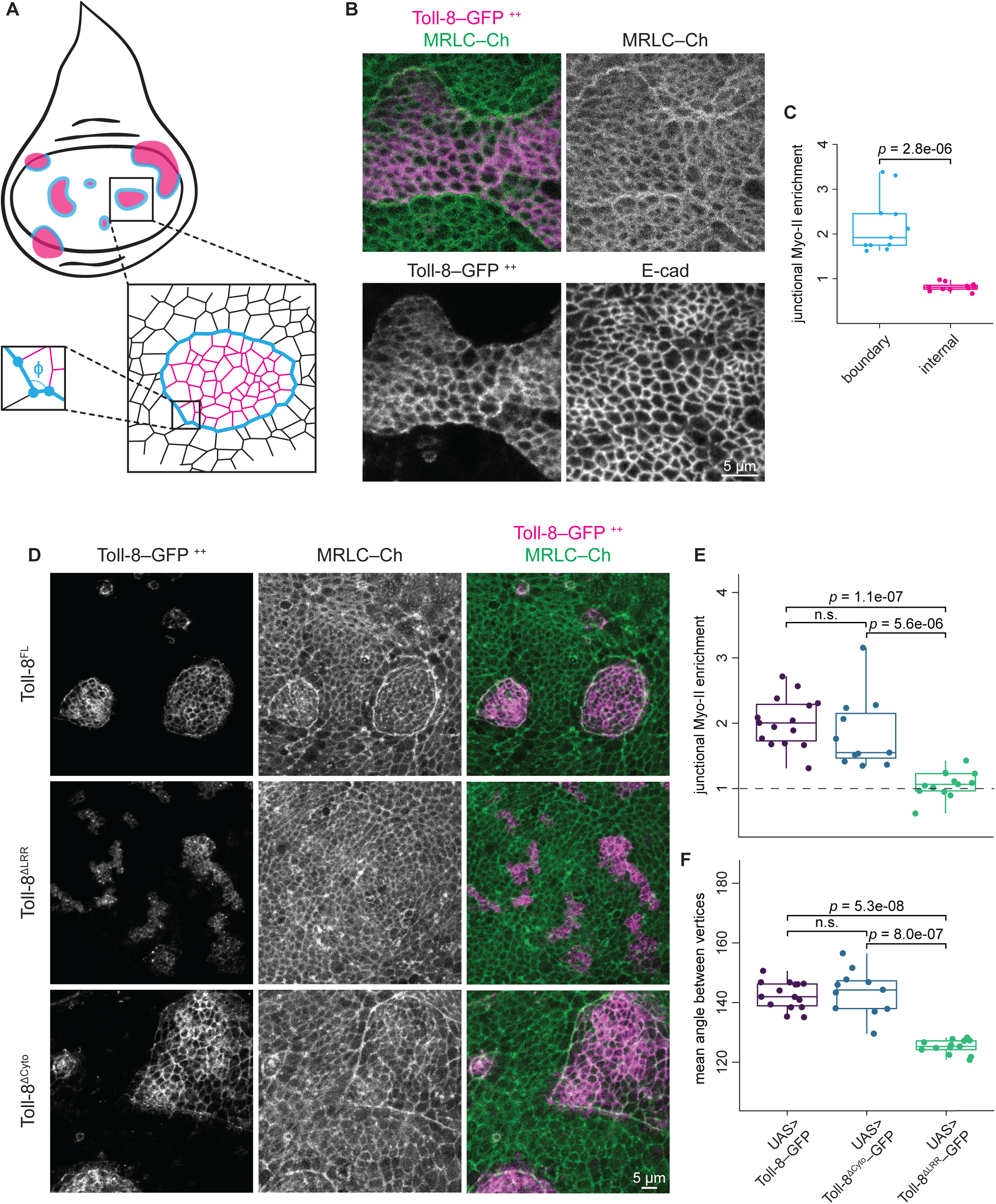
Toll-8 asymmetry leads to Myo-II enrichment in wing discs, independent of the Toll-8 cytoplasmic tail. **(A)** Schema showing wing disc clones (pink, top) and a detailed view of cell interfaces around a clone (bottom). Interfaces within the clone in pink; those within the wild-type tissue in black; those at the clonal boundary in blue. ϕ is the angle between neighboring vertices at the clonal boundary. **(B)** Fixed Toll-8–GFP (bottom left), MRLC–Ch (top right) and E-cad (bottom right) signals from wing disc clones overexpressing full length Toll-8 (*n*=11). Myo-II is enriched at the boundary of the clone but not inside the clone. **(C)** Junctional Myo-II enrichment relative to wild-type tissue within Toll-8 overexpressing clones (pink) or at the boundary of Toll-8 overexpressing clones (blue) for the condition shown in (**B**). **(D)** Fixed Toll-8–GFP (left) and MRLC–Ch (middle) signals from wing disc clones overexpressing full length Toll-8 (Toll-8^FL^, top, *n*=15), Toll-8 with the extracellular LRRs removed (Toll-8^ΔLRR^, middle, *n*=11), or Toll-8 with the intracellular cytoplasmic tail removed (Toll-8^ΔCyto^, bottom, *n*=13). Myo-II enrichment requires the extracellular LRRs but not the cytoplasmic tail of Toll-8. **(E and F)** Junctional Myo-II enrichment (**E**) and boundary smoothness (**F**) are quantified at clone boundaries for the conditions shown in (**D**). Scale bars: 5 µm. Statistics: Mann-Whitney U test; n.s.: *p* > 0.05.

The cytoplasmic domain of Toll proteins is necessary for canonical Toll signal transduction (Anthoney et al., 2018). To test whether the cytoplasmic domain of Toll-8 is required for Myo-II enrichment, we generated wing disc clones overexpressing a truncated version of Toll-8, removing its cytoplasmic domain (Toll-8^ΔCyto^). Surprisingly, Myo-II was still enriched at the boundary of Toll-8^ΔCyto^ overexpressing clones (Figures 2D and 2E), and the clonal boundary was still smooth compared to Toll-8 ^ΔLRR^ clones (Figures 2D and 2F). We conclude that the cytoplasmic domain of Toll-8 is dispensable for Myo-II polarization. We observed similar results for Toll-6 and Toll-2, which also enrich Myo-II at clonal boundary independent of their cytoplasmic tails (Figures S2A and S2B). This indicates that the signaling events leading to Myo-II enrichment at the boundary between Toll-8-expressing and non-expressing cells may require another protein, presumably interacting with Toll-8 at the cell surface.

### Cirl/Latrophilin binds Toll-8 and mediates Toll-8-induced Myo-II enrichment

To look for binding partners of Toll-8, we performed affinity purification-mass spectrometry experiments using lysates isolated from embryos overexpressing Toll-8–YFP as a bait (Figure 3A and Methods). Of the specifically bound proteins, Cirl was the most abundant target (Figure 3B and Table 1) and of particular interest since it is a G protein-coupled receptor (GPCR). Cirl is the *Drosophila* homologue of vertebrate Latrophilin, a member of the adhesion GPCR (aGPCR) subfamily (Scholz et al., 2015; Schöneberg and Prömel, 2019). Interestingly, the Toll-8 extracellular domain shares sequence similarities with the extracellular domains of the human LRR proteins FLRTs (Dolan et al., 2007), which are known to form protein complexes with human Latrophilins (Boucard et al., 2014; O’Sullivan et al., 2012; Sando et al., 2019; del Toro et al., 2020). We also recovered Toll-2 as a significant Toll-8 binding partner *in vivo* (Figure 3B and Table 1), in agreement with previous findings *in vitro* (Paré et al., 2014).

**Figure 3.**
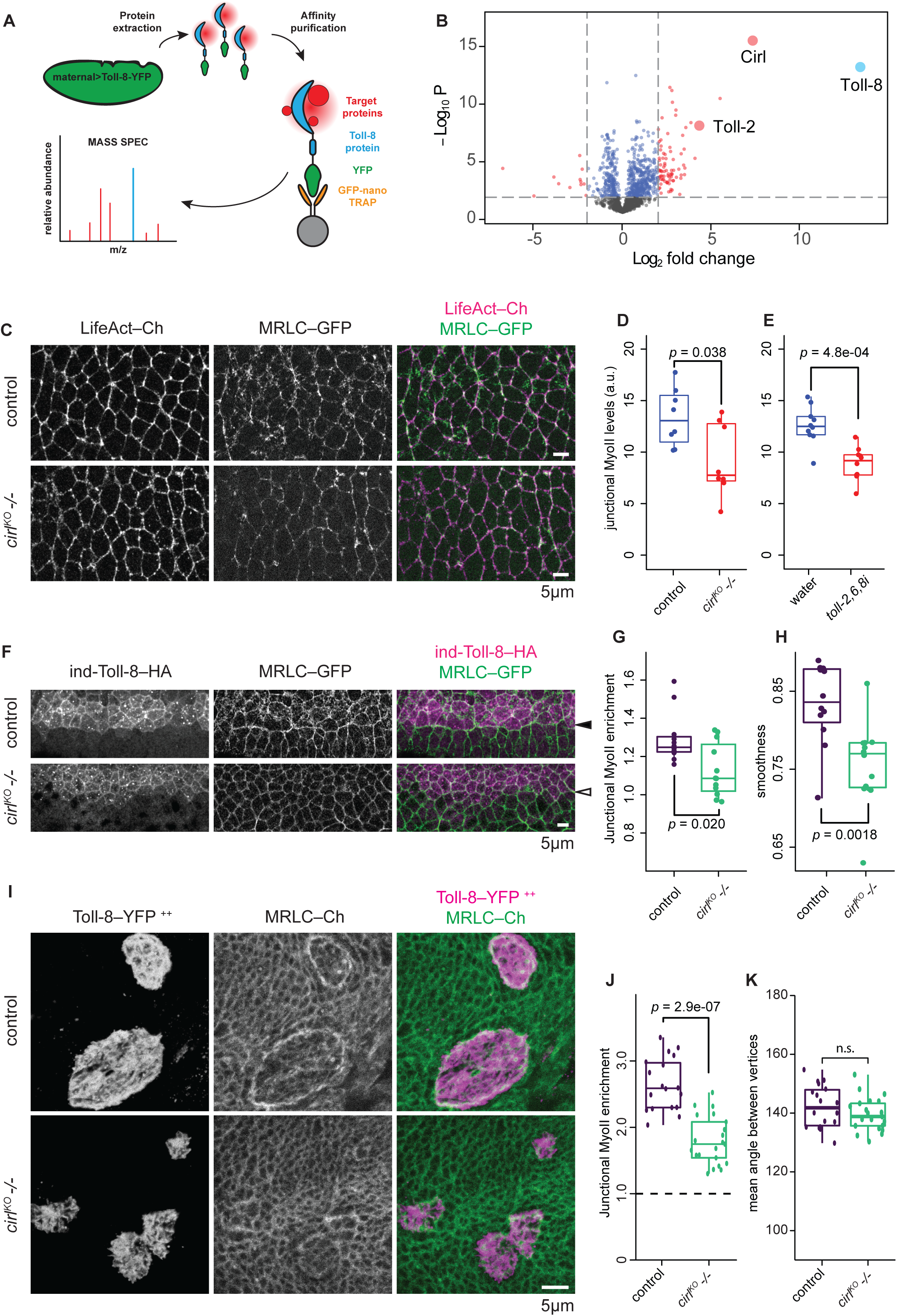
The adhesion GPCR Cirl interacts physically with Toll-8 and is necessary for Myo-II enrichment induced by Toll-8. **(A)** Schematics showing the procedure of affinity purification-mass spectrometry using embryos overexpressing the Toll-8–YFP fusion protein (Toll-8 protein in blue, YFP tag in green, potential binding targets in red) and the GFP-Trap (nanobody against GFP/YFP in orange, agarose bead in grey). See methods for details. **(B)** Volcano plot showing differential levels (Log_2_|fold change|, x axis) and p-values (-Log_10_ P, y axis) of proteins identified by mass spectrometry in *Drosophila* embryos overexpressing Toll-8–YFP versus *wt* embryos. Vertical dashed lines denote the cut-off for log2|Fold Change| as 2. Horizontal dashed line denotes the cut-off for p-value as 0.01. Toll-8 (bait), Cirl and Toll-2 are highlighted. **(C)** Still images from time lapse movies in *wt* (top, *n*=8) or *cirl^KO^ -/-* (bottom, *n*=8) embryos. LifeAct– Ch marks cell outlines. Junctional Myo-II is reduced in *cirl^KO^ -/-* null mutant embryos. **(D)** Quantifications of mean junctional Myo-II levels for the conditions shown in (**C**). **(E)** Quantifications of mean junctional Myo-II levels in embryos injected with water (*n*=10) or dsRNAs against Toll-2,6,8 (*n*=10) (related to Figures S1A and S1B). **(F)** Anti-Toll-8–HA (left) and anti-MRLC–GFP (middle) signals in *wt* (top, *n*=12) or *cirl^KO^ -/-* (bottom, *n*=12) embryos expressing full length Toll-8 under the *ind* promoter. Myo-II is reduced at the *ind* ventral boundary in *cirl^KO^ -/-* embryos (empty arrowhead). **(G and H)** Junctional Myo-II enrichment (**G**) and boundary smoothness (**H**) at the ventral *ind* boundary for the conditions shown in (**F**). **(I)** Fixed Toll-8–YFP (left) and MRLC–Ch (middle) signals from wing disc clones overexpressing full length Toll-8 in *wt* (top, *n*=18) or *cirl^KO^ -/-* (bottom, *n*=21) wing discs. Myo-II is reduced at the boundary of Toll-8 overexpressing clones in *cirl^KO^ -/-* wing discs. **(J and K)** Junctional Myo-II enrichment (**J**) and boundary smoothness (**K**) at clone boundaries for the conditions shown in (**I**). Scale bars: 5 µm. Statistics: Mann-Whitney U test; n.s.: *p* > 0.05.

**Table 1:** List of proteins associated with Toll-8 (in green) selected using pFDR at 0.1% and s0=1. The proteomes of embryos overexpressing Toll-8–YFP maternally and zygotically (N = 3 protein isolations: T3, T4, T5, each with 3 replicas: T3a, T3b, T3c, …) were compared with yw embryos (N = 3 protein isolations: C3, C4, C5, each with 3 replicas: C3a, C3b, C3c, …) with affinity purification mass spectrometry (PRIDE dataset identifier PXD017895). Protein list is displayed by order of Difference log2(T/C). Rows highlighted in green denote significant hits from a two-sample t-test using permutation-based false discovery rate (pFDR) controlled at 0.1% (5000 permutations).

Using a Cirl–RFP knock-in line (Scholz et al., 2017) to visualize endogenous Cirl, we found that Cirl localized to the membrane and was enriched at cell-cell interfaces around adherens junctions in both the embryo and the wing disc (Figures S3A and S3B). We next asked whether Cirl is required for Myo-II enrichment in the embryo. *cirl* maternal and zygotic mutant embryos showed delayed extension of the embryonic axis (Figures S3C and S3D), reduced cell intercalation (Figures S3E and S3F; Movie S1), and a strong reduction of junctional Myo-II in the ectoderm (Figures 3C and 3D), resembling Myo-II reduction observed in *toll-2,6,8* RNAi embryos (Figures 3E and S1A). Thus, *cirl* is required for junctional Myo-II enrichment in the embryo.

We further tested whether Cirl is required for Toll-8-induced polarization of Myo-II by comparing the ventral boundary of *ind-Toll-8–HA* stripe in wild-type and *cirl* null mutant embryos. In the absence of Cirl, we found a strong reduction of Myo-II enrichment at the boundary of Toll-8 stripe, which was also less smooth (Figures 3F-3H). Moreover, in wing imaginal discs we observed a similar reduction in Myo-II enrichment at the boundary of clones overexpressing Toll-8–YFP in a *cirl* null mutant background (Figures 3I and 3J). Note that Myo-II enrichment was not completely abolished and the clonal boundary was still smooth (Figure 3K), suggesting that other molecules in addition to Cirl may interact with Toll-8 to confer Myo-II enrichment at clone boundaries in the wing disc. Taken together, we conclude that Toll-8 and Cirl physically and functionally interact to polarize Myo-II at the interface between Toll-8 expressing and non-expressing cells.

### Cirl asymmetric interfaces lead to Myo-II enrichment

Interestingly, we observed that Myo-II was enriched on both sides of the interface between Toll-8 expressing and non-expressing cells (Figure S4A, cyan arrowheads), indicating that Myo-II is also enriched in cells that do not express Toll-8. We thus tested whether Cirl mediates Myo-II enrichment on both sides of the interface. To this end, we performed mosaic analysis with a repressible cell marker (MARCM) in wing discs and induced *cirl* null mutant clones either adjacent to (Figure 4A) or coincident with (Figure 4B) Toll-8 overexpressing clones. We expected that removing Cirl from one side should lead to a reduction in Myo-II enrichment at the clone boundary due to a loss of Myo-II enrichment in cells where Cirl is absent. Surprisingly, when Cirl was only absent from neighboring cells, Myo-II was enriched at the Toll-8 boundary at similar levels to control interfaces (Figure 4A, compare orange and cyan arrows, quantified in Figure 4C) and on both sides of the clone boundary (Figure S4A). Similarly, when Cirl was only absent from cells overexpressing Toll-8, Myo-II was still enriched at similar levels at the clone boundary (Figure 4B, quantified in Figure 4C) and on both sides of the clone boundary (Figure S4B). Thus, Cirl is dispensable in either Toll-8 expressing or responding/contacting cells. This is remarkable since Cirl must be present on at least one side of Toll-8 overexpressing clones, as the complete removal of Cirl on both sides significantly reduces Myo-II enrichment (Figure 3J). We thus hypothesized that 1) the role of Toll-8 is to induce an asymmetry of Cirl activity at the clonal boundary, and that 2) Cirl asymmetric interfaces enrich Myo-II. To test the latter, we generated *cirl* mutant clones in the wing disc without overexpressing Toll-8. We reasoned that at the *cirl* mutant clonal interface Cirl localization and activity is *de facto* asymmetric. This is supported by the observation that Cirl is still localized in wild-type cell interfaces in contact with *cirl* mutant cells (Figure S4C, yellow arrowhead). We found that Myo-II was indeed enriched at the boundary of *cirl* mutant clones and that the boundary was smooth compared to control clones (Figures 4D-4F). This is similar to Toll-8 overexpressing clones, albeit to a lesser extent. We thus conclude that asymmetric Cirl interfaces at *cirl* mutant clonal boundary enrich Myo-II. Hence, we next investigated whether Toll-8 does induce a Cirl asymmetry at its expression boundary.

**Figure 4.**
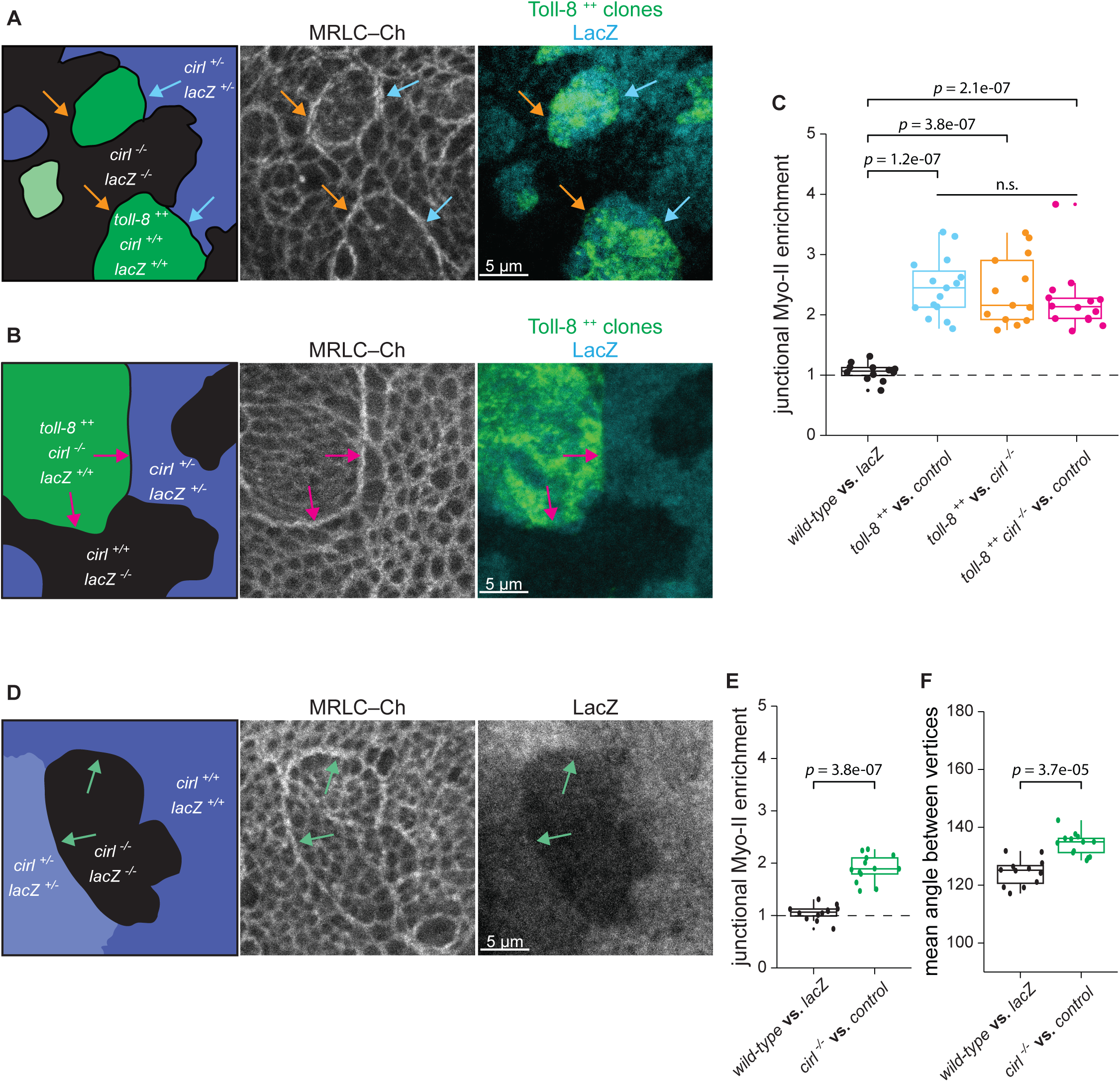
Cirl asymmetric expression leads to Myo-II enrichment. **(A)** MARCM clones in a wing disc where clones overexpressing Toll-8–YFP (*toll-8 ^++^* in green) are juxtaposed to control cells heterozygous (*cirl ^+/-^* in blue, cyan arrows, *n*=15) or null mutant (*cirl ^-/-^* in black, orange arrows, *n*=13) for *cirl*. Myo-II enrichment at the boundary of Toll-8 overexpressing cells is similar in both cases. **(B)** MARCM clones in the wing disc. Myo-II is enriched at boundaries (magenta arrows, *n*=14) of clones overexpressing Toll-8– YFP and null mutant for *cirl* (*toll-8 ^++^, cirl ^-/-^* in green) juxtaposed to cells heterozygous (*cirl ^+/-^* in blue) or *wild-type* (*cirl ^+/+^* in black) for *cirl*. **(C)** Quantifications of junctional Myo-II enrichment at clone boundaries for the conditions shown in (**A**) and (**B**). With the exception of control clones (black boxplot, images not shown, *n*=12), colors of the boxplots correspond to the arrows in (**A**) and (**B**). **(D)** Myo-II is enriched at the boundary (green arrows) of *cirl^-/-^* null mutant clones (*cirl^-/-^* in black, *n*=13) in the wing disc. **(E and F)** Junctional Myo-II enrichment (**E**) and boundary smoothness (**F**) quantified at clone boundaries for the conditions shown in (**D**). The black plot (images not shown) in (**E**) is the same as in (**C**). Scale bars: 5 µm. Statistics: Mann-Whitney U test; n.s.: *p* > 0.05.

### Toll-8 generates a Cirl interfacial asymmetry at its expression boundary

To test if Toll-8 is able to induce a Cirl asymmetry at its expression boundary, we first assessed the effect of Toll-8 overexpressing clones on Cirl localization with wing discs expressing Cirl– RFP from the endogenous locus. In control cells that do not overexpress Toll-8, Cirl–RFP was localized at adherens junctions (marked by E-cadherin localization) and in the subapical domain (i.e. the domain above adherens junctions where neighboring cells are in direct contact) (Figures 5A-5A’’, black curve). In cells overexpressing Toll-8–YFP, Cirl–RFP surface levels increased both at adherens junctions and in the subapical domain (Figures 5A-5A’’, magenta curve shifted compared to black curve). Cirl–RFP distribution follows that of Toll-8–YFP (Figures 5A, 5A’, see overlays, and 5A’’’), suggesting that Toll-8 re-localizes and stabilizes Cirl. At high levels of expression, Toll-8–YFP was present at the free apical membrane (i.e. the domain where cells are in contact with the extracellular space), which coincided with ectopic localization of Cirl–RFP in this domain (Figure 5A’, yellow arrowheads). This suggests that Toll-8 and Cirl physically interact *in cis*. Strikingly, we observed that Cirl–RFP was depleted from the orthogonal junctions in wild-type cells in direct contact with the boundary of Toll-8–YFP overexpressing cells (Figures 5A, yellow arrow and S5A). Thus, Toll-8 affects Cirl localization *in trans*. We observed similar effects on Cirl localization with Toll-6 clonal overexpression (Figure S5B) but not with Toll-2 (Figure S5C), suggesting that Toll-6 also interacts with Cirl while Toll-2 might interact with another protein.

**Figure 5.**
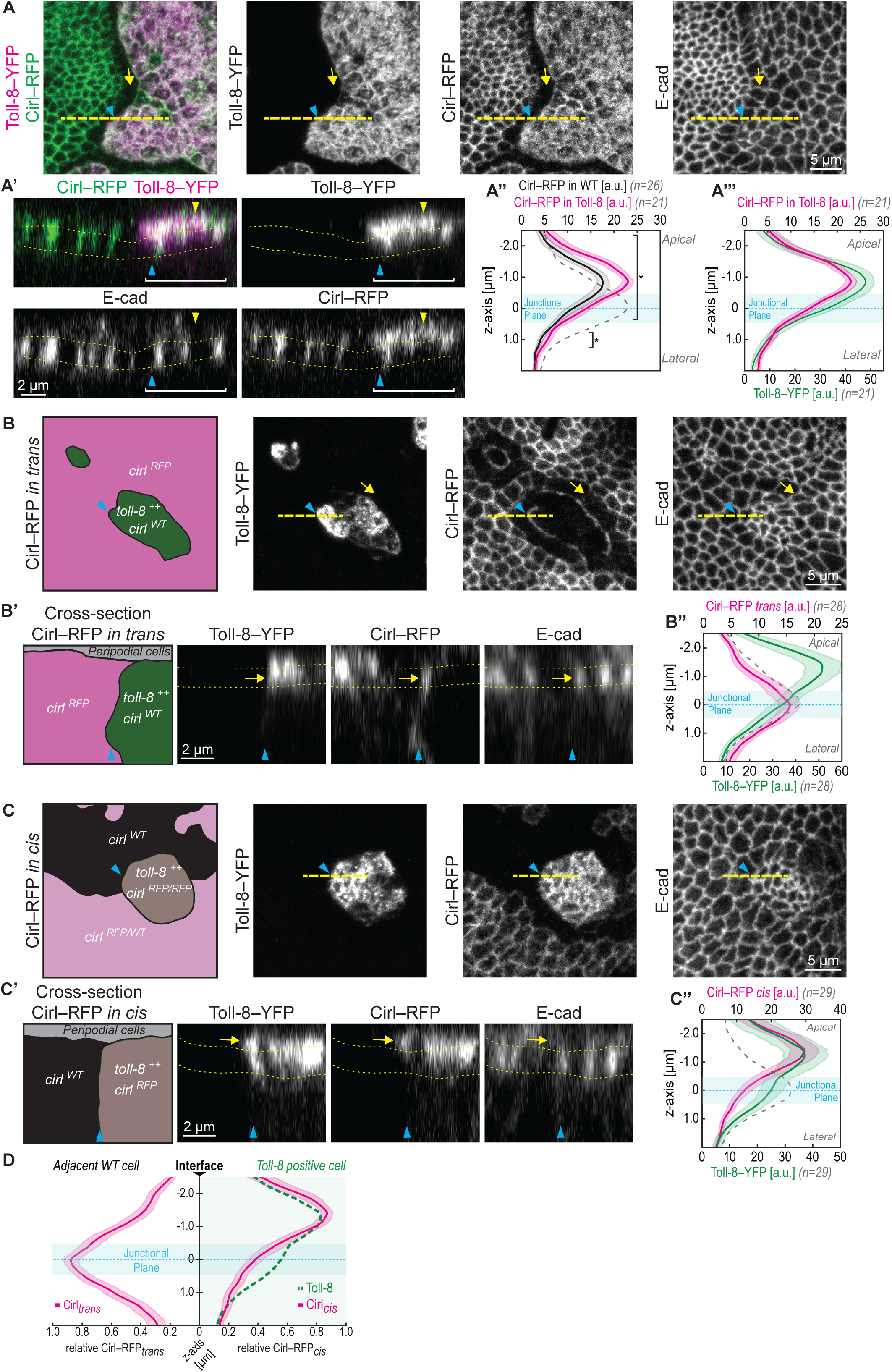
Toll-8 generates Cirl interfacial asymmetry at the boundary of its expression domain. Blue arrowheads indicate clone boundaries in all panels. **(A)** Toll-8–YFP overexpressing clone in a Cirl–RFP wing disc. Toll-8–YFP (magenta) and anti Cirl–RFP (green) signals colocalize inside the clone. Cirl–RFP is depleted from junctions orthogonal to the clone boundary (yellow arrow). **(A’)** Optical cross-section of the position indicated by the dashed line in (**A**) (apical to the top). Dashed lines in (**A’**) mark the position of the junctional plane based on E-cad signals. Yellow arrowhead shows presence of both Toll-8 and Cirl at the free apical membrane. **(A’’ and A’’’)** Intensity quantifications (x-axis) along the apicobasal axis (y-axis) for Toll-8 and Cirl. The junctional plane (blue area) is defined by E-cad peak levels (dashed grey curve). Cirl–RFP levels increase in Toll-8 overexpressing clones both at the junctional plane and in the subapical domain (**A’’**, square brackets) following Toll-8 distribution (**A’’’**). **(B)** Toll-8–YFP overexpressing clones (*toll-8 ^++^*, *cirl ^WT^* in green) in a wing disc where Cirl–RFP is present only outside of Toll-8 overexpressing clones (*cirl ^RFP^* in magenta). Cirl–RFP is depleted from interfaces orthogonal to the clone boundary (yellow arrow) and accumulates at the clone boundary (blue arrowhead). **(B’)** Optical cross-section (dashed line in (**B**)) shows Cirl–RFP enrichment at the junctional plane (yellow arrow), quantified in (**B’’**). **(C)** Toll-8–YFP overexpressing clone with Cirl–RFP present inside (*toll- 8 ^++^*, *cirl ^RFP^* in brown) facing cells homozygous for untagged Cirl (*cirl ^WT^* in black) or heterozygous for *cirl ^RFP^* (*cirl ^RFP/WT^* in light magenta). **(C’)** Optical cross-section (dashed line in (**C**)) shows Cirl–RFP enrichment above the junctional plane (yellow arrow), quantified in (**C’’**). **(D)** Comparison of Cirl–RFP apicobasal localization at both sides of Toll-8 clonal boundaries. Cirl is observed at the junctional plane *in trans* and above the junctional plane *in cis* creating a Cirl interfacial asymmetry at Toll-8 clonal boundaries. Scale bars: 5 µm in (**A**), **(B)** and (**C**); 2 µm in (**A’**), (**B’**) and (**C’**). Error bands indicate the 95% confidence interval in all plots. Statistics in (**A’’**): two-sided t-test; *: *p* < 0.05.

The depletion of Cirl from orthogonal junctions in contact with Toll-8 overexpressing cells led us to ask if Cirl is polarized at the boundary of Toll-8 overexpressing cells *in trans*. To test this, we used MARCM to observe endogenously tagged Cirl–RFP only adjacent to (i.e. *in trans*) Toll-8–YFP overexpressing cells. Note that in Cirl–RFP negative cells, untagged endogenous Cirl is present. When Cirl–RFP was present only *in trans*, Cirl–RFP was indeed localized at the clone boundary (Figure 5B, blue arrowheads) and depleted from the junctions orthogonal to the clone boundary (Figure 5B, yellow arrows). This effect did not require the presence of Cirl inside Toll-8 overexpressing cells (Figure S5D). Thus, Cirl is planar polarized in wild-type cells in direct contact with Toll-8 overexpressing cells, suggesting that Toll-8 and Cirl physically interact *in trans*. Surprisingly, at the clone boundary, Cirl–RFP *in trans* strictly colocalized with E-cad at adherens junctions and was absent from the subapical domain (Figures 5B’, arrow, and 5B’’). We then examined how Cirl–RFP was localized at clonal boundaries in cells overexpressing Toll-8–YFP using MARCM to observe endogenously tagged Cirl–RFP only inside the clone (i.e. *in cis*). When Cirl–RFP was present only *in cis* (Figure 5C), Cirl–RFP was enriched in the subapical domain and tended to be reduced at adherens junctions (Figures 5C’, arrow, and 5C’’). Therefore, at cell-cell interfaces between wild-type and Toll-8 overexpressing cells, where cells are in direct contact, Cirl is specifically localized to adherens junctions *in trans* (Figures 5B’ and 5B’’) while it is enriched above adherens junctions *in cis* (Figures 5C’ and 5C’’), creating an interfacial asymmetry in Cirl localization (Figure 5D).

Toll-8 can thus induce an interfacial asymmetry in Cirl apicobasal localization at Toll-8 expression boundary. Our genetic setup does not allow us to assess Cirl localization only at one side of cell interfaces inside Toll-8 overexpressing clones. Given that inside the clone Toll-8 is symmetrically present both *in cis* and *in trans*, we assume that Cirl should be symmetrically localized at these internal interfaces, resulting in Cirl interfacial asymmetry specifically at the Toll-8 expression boundary, and not at internal interfaces. Since we showed that asymmetric Cirl interfaces between wild-type and *cirl* mutant cells lead to Myo-II enrichment (Figures 4D and 4E), we propose that the interfacial asymmetry of Cirl induced by Toll-8 might be a signal leading to Myo-II enrichment at Toll-8 expression boundaries.

### Quantitative differences in Toll-8 expression lead to mutually dependent Toll-8 and Cirl polarity

To gain further insights into the dynamic process of Myo-II planar polarization, we performed *ex vivo* live imaging (Dye et al., 2017) of nascent Toll-8 overexpressing clones in cultured wing discs. We used a temperature-sensitive GAL80 (GAL80ts) to precisely time the onset of Toll-8 expression by a temperature shift to 30°C. After 2h15 at 30°, we isolated wing discs with Toll-8–YFP clones and expressing MRLC–mCherry and imaged them in a thermostatic chamber at 30°C. This allowed us to analyze Myo-II planar polarization in response to Toll-8 levels, which were low at the beginning and increased over time. Myo-II enrichment was already observed when Toll-8–YFP was initially restricted to cell-cell interfaces, prior to its subsequent accumulation in the free apical membrane (Figure S6A). This supports the idea that interfacial Toll-8–YFP, instead of Toll-8 in the free apical membrane, is responsible for polarization of Myo-II.

We took advantage of the fact that in this experimental setup Toll-8 expression was not induced synchronously in all cells of a given clone, likely due to stochasticity in GAL80ts inactivation/GAL4 de-repression (Figure 6A and Movies S2-S4). This led to the generation of dynamic changes in Toll-8 expression creating quantitative differences in Toll-8 expression levels between neighboring cells. This opened the possibility to correlate quantitative differences in Toll-8 expression and Myo-II planar polarization. We found that Myo-II was not only enriched at the boundary of Toll-8–YFP expressing cells facing Toll-8–YFP negative cells as in previous experiments, but also enriched at interfaces between cells with different Toll-8– YFP levels (Figures 6A, 110’, cyan arrow, and S6B; Movies S2 and S3). In this assay, the kinetics of Myo-II polarization at the boundary of cells expressing different levels of Toll-8 is in the range of 10 min (Figure 6A’, between 40 min and 50 min, Myo-II becomes polarized at a Toll-8 interface) and the amplitude of Myo-II polarity is around two fold (Figure S6B), which is commensurate with the dynamics and amplitude of Myo-II polarization in the embryonic ectoderm (Bertet et al., 2004; Blankenship et al., 2006). Moreover, as the levels of Toll-8–YFP further increased, and once Toll-8–YFP expression reached the same level between these contacting cells, Myo-II enrichment was no longer present at these interfaces (Figure 6A’, arrowheads), and was stabilized only at the boundary of the clone (Figures 6A, 330min, and S6B). This argues that quantitative differences in Toll-8 expression between neighboring cells polarize Myo-II.

**Figure 6.**
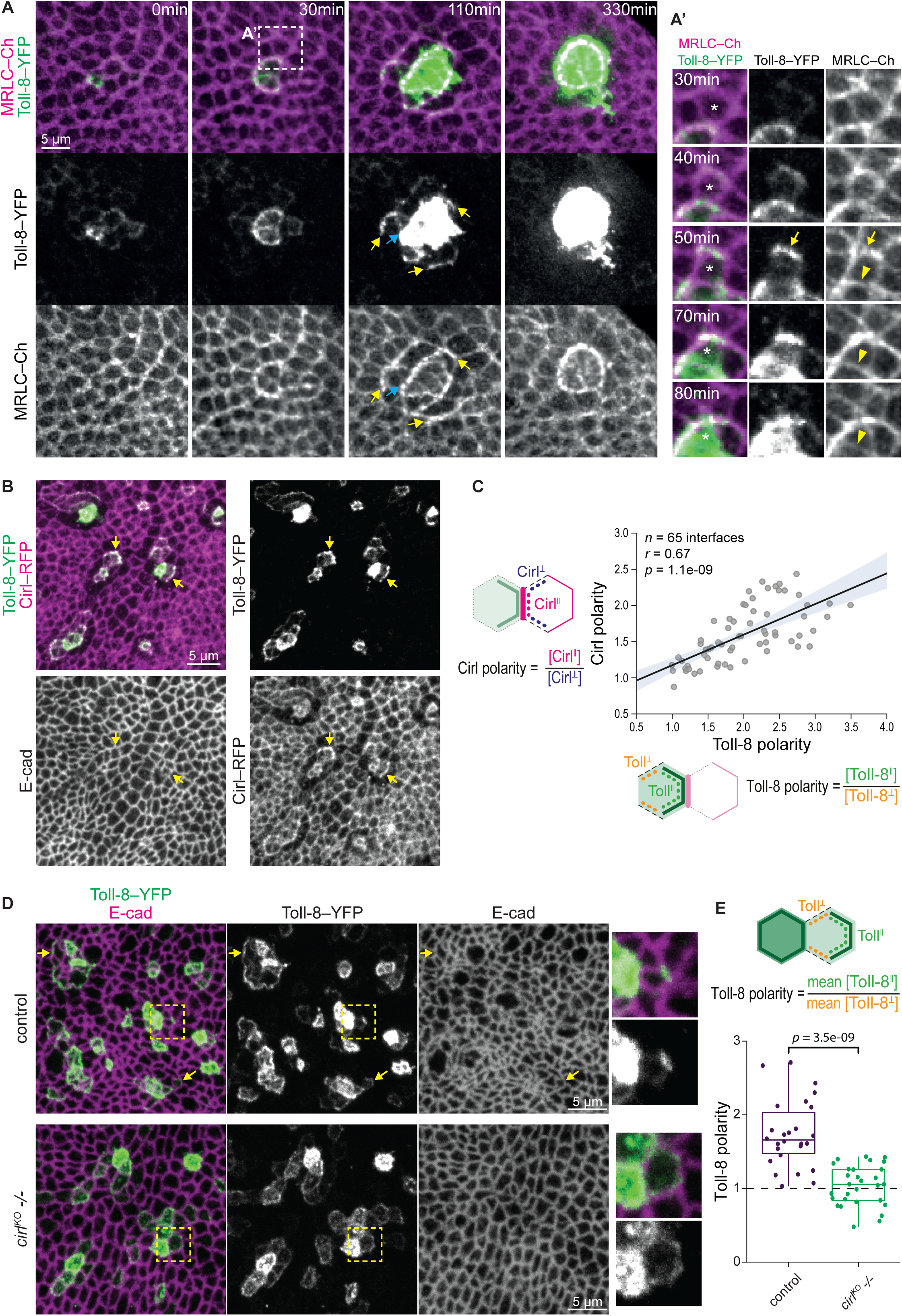
Quantitative differences in Toll-8 expression lead to mutually dependent Toll- 8 and Cirl polarity. **(A)** Stills from a time lapse movie of a wing disc showing live dynamics of Toll-8–YFP (green) and MRLC–Ch (magenta) in a nascent Toll-8–YFP overexpressing clone. Toll-8– YFP starts being detectable at 0 min and its levels increase during the course of the time lapse. Yellow arrows mark junctions displaying Toll-8 planar polarized enrichment associated with Myo-II enrichment. Cyan arrow indicates an interface between high and low Toll-8 overexpressing cells. **(A’)** Zoomed view of a cell (asterisk) that dynamically upregulates expression of Toll-8–YFP during ∼50min. Myo-II is being enriched at junctions that show Toll-8–YFP planar polarity (arrow) and this enrichment is lost once Toll-8–YFP expression reaches similar levels between contacting cells (arrowhead). **(B)** Nascent Toll-8– YFP clones (green) in a Cirl–RFP (magenta) wing disc. Cirl levels are increased at Toll-8– YFP enriched junctions (arrows). **(C)** Quantifications of Toll-8–YFP planar polarity versus Cirl–RFP planar polarity *in trans* for the condition shown in (**B**) indicate a significant positive correlation between the two variables. **(D)** Toll-8–YFP planar polarity in nascent Toll-8–YFP clones in control (top, *n*=26) or *cirl* null mutant (bottom, *n*=29) wing discs. Zoomed views are shown on the right. Toll-8–YFP planar polarity is reduced in the absence of Cirl. **(E)** Quantifications of Toll-8–YFP planar polarity in cells expressing low Toll-8– YFP levels in direct contact with cells expressing high Toll-8–YFP levels for the conditions shown in (**D**). Scale bars: 5 µm. Error band in (**C**) indicates the 95% confidence interval. Statistics in (**E**): Mann-Whitney U test.

Strikingly, between neighboring cells with quantitative differences in Toll-8 expression levels, Toll-8–YFP was initially planar polarized in cells expressing lower levels and tended to accumulate at interfaces facing away from the cells expressing higher levels (Figures 6A and A’, yellow arrows; Movies S2-S4). Moreover, in cells where Toll-8–YFP was planar polarized, Myo-II was specifically enriched at Toll-8-enriched interfaces, leading to Myo-II planar polarity across several rows of cells (Figure 6A, 110min, yellow arrows; Movies S2 and S3). Therefore, Toll-8 polarization emerged when neighboring cells express different levels of Toll-8 and this is correlated with polarization of Myo-II.

We then asked how Cirl localization was affected by these nascent polarized patterns of Toll-8. Since Cirl–RFP was too low to be detected in our live imaging setup, we fixed wing discs treated under the same conditions as above (see Methods). We found that Cirl–RFP was enriched at interfaces where Toll-8 was planar polarized (Figure 6B, arrows). In addition, Cirl was reduced in orthogonal interfaces facing cells exhibiting Toll-8 planar polarity and we found a significant correlation between Toll-8 planar polarity and Cirl planar polarity in the neighboring cells *in trans* (Figures 6B and 6C). Therefore, we conclude that Cirl forms planar polarized patterns in response to Toll-8 polarization. Note that in this experiment, we could not assess Cirl interfacial asymmetry as done in Figure 5 due to the complexity of the genetics required.

Finally, we asked if this transient Toll-8 planar polarity requires Cirl. To test this, we induced nascent Toll-8 overexpression clones in wild-type or *cirl* mutant wing discs. We found that Toll-8 planar polarity was strongly reduced between cells expressing different levels of Toll-8 in the absence of Cirl (Figure 6D and 6E). Thus, the planar polarity of Toll-8 and Cirl mutually depend upon one another. Since Toll-8 and Cirl form a molecular complex, we propose that Toll-8 and Cirl mutually attract each other in a positive feedback, which results in their self-organized polarity at cell interfaces.

## Discussion

Our work sheds new light on the question of how Myosin-II planar polarity emerges from the juxtaposition of cells with distinct genetic identities. We report that Toll-8 asymmetric expression leads to Myo-II polarization independent of other Toll receptors. We identified a cell surface protein complex between Toll-8 and the adhesion GPCR Cirl/Latrophilin that is required for Myo-II polarization. Though complete removal of Cirl diminishes Toll-8-induced polarization of Myo-II, removing Cirl from either side of a Toll-8 expression asymmetric interface does not affect Myo-II polarization. This suggests that Toll-8 induces Cirl asymmetry at Toll-8 expression boundary and that this asymmetry itself leads to Myo-II activation. Consistent with this, Cirl asymmetric expression alone enriches Myosin-II. Moreover, we observed that at the boundary of Toll-8 expression domain, Cirl is differentially localized *in trans* and *in cis* thus creating a Cirl interfacial asymmetry. This suggests that Cirl asymmetry could be a potential signal for Myo-II enrichment. We have shown that Toll-8/Cirl co-polarity emerges from quantitative differences in Toll-8 expression between neighboring cells, via a mutual positive feedback mechanism. Thus, we propose a conceptual model wherein Toll-8 and Cirl generate self-organized polarity, which leads to Myo-II enrichment (Figure 7): the first cell expressing Toll-8 (cell A) recruits Cirl *in trans* from neighboring cells by depleting it from their orthogonal interfaces (Figure 7, dashed lines), resulting in Cirl planar polarity in these neighboring cells (Figure 7, panel 1). When one of these cells (cell B) initiates expression of Toll-8, Toll-8 expression levels are lower than in cell A. Due to this quantitative difference in Toll-8 expression and a preexisting Cirl planar polarity in cell B (where Cirl is depleted from orthogonal interfaces in contact with cell A), Toll-8 in cell B is attracted to the remaining interfaces containing Cirl *in trans*, which are facing away from cell A (Figure 7, panel 2, green arrows and green line in cell B). These new Toll-8-enriched interfaces will stabilize even more Cirl *in trans* at the expense of orthogonal interfaces in the cells adjacent to cell B that do not express Toll-8, thus propagating Cirl planar polarity one row of cells further away (Figure 7, panel 2, magenta arrows). In summary, we propose that quantitative differences in Toll-8 expression between neighboring cells translate into self-organized Toll-8/Cirl/Myo-II planar polarity due to local interactions between Toll-8 and Cirl.

**Figure 7.**
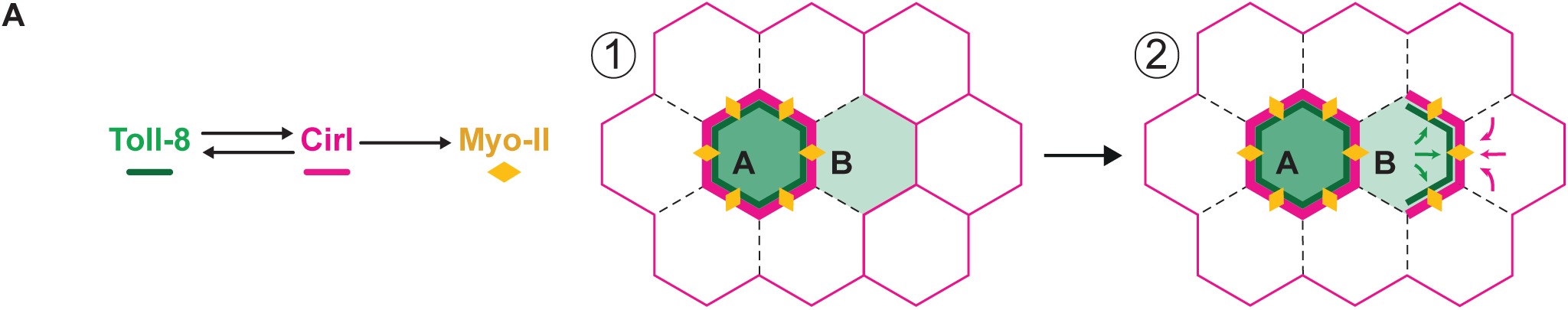
Toll-8 and Cirl form dynamic self-organized patterns. **(A)** Model proposing mutual attraction between Toll-8 and Cirl leading to the formation of self-organized planar polarized patterns of Toll-8/Cirl/Myo-II (refer to the discussion for a detailed description). Dashed lines represent interfaces depleted of Cirl.

We propose that Toll-8 enriches Myo-II through the asymmetric localization of Cirl. The downstream signaling pathway linking interfacial asymmetry of Cirl to Myo-II on both sides of the Toll-8 expression boundary remains to be elucidated. What may prevent symmetric Cirl from signaling to enrich Myo-II? Contrary to the adhesion GPCR Flamingo/CELSR (Usui et al., 1999), Cirl does not require homophilic interaction *in trans* to localize at the membrane, but interestingly in *C. elegans* Latrophilin extracellular domains form stable dimers (Prömel et al., 2012). Hence, extracellular dimerization of Cirl *in trans* could be a mechanism inhibiting Cirl signaling when it is localized symmetrically between neighboring cells. The asymmetric localization of Cirl induced at the boundary of Toll-8 expression could dissociate the Cirl dimers, release auto-inhibition and thereby elicit signaling.

The fact that Myo-II enrichment happens on both sides of the Toll-8 clonal boundaries, even in the absence of Cirl on one side of the boundary, is intriguing. The asymmetry of Cirl extracellular domains may activate other transmembrane proteins present on both sides of the clone boundary, in particular other GPCRs such as Smog (Kerridge et al., 2016), which then signals bidirectionally to enrich Myo-II. Alternatively, Cirl asymmetry may elicit Myo-II polarity by differential recruitment of Rho1 regulators, akin to Cadherin2 in *Ciona* (Hashimoto and Munro, 2019), enriching Myo-II through its intracellular domain on one side of the interface and propagating Myo-II enrichment on the other side of the interface through mechanical feedback.

It was proposed that heterophilic interactions between Toll receptors were required to recruit Myo-II (Paré et al., 2014). Here we show that asymmetric expression of a single Toll receptor, Toll-8, enriches Myo-II independent of other Toll receptors through feedback interactions with the adhesion GPCR Cirl/Latrophilin. Although clonal analysis is not possible in early *Drosophila* embryos, preventing the dissection of how a Cirl interfacial asymmetry might be generated in this tissue, we propose two hypotheses on how several Toll receptors and Cirl may interact to define planar polarized interfaces for Myo-II enrichment in this tissue. On the one hand, as Toll-2, 6 and 8 all have distinct expression patterns and boundaries in the embryo (Paré et al., 2014), and since Toll-8 alone is capable of polarizing Myo-II in embryos and discs, local Cirl asymmetry could be induced at each boundary independently, thereby allowing polarization on Myo-II at all interfaces on the basis of what we report here with Toll-8/Cirl (Tetley et al., 2016). On the other hand, interactions between different Toll receptors (Paré et al., 2014 and Table 1) might modulate Toll-8/Cirl binding. This could lead to Cirl asymmetry at vertical interfaces across which different combinations of Toll receptors are expressed, contrary to horizontal interfaces that expressed similar combination of Toll receptors on both sides.

We observed in the wing disc that various quantitative and spatial patterns of Toll-8 have different outputs. Groups of cells expressing homogeneously high levels of Toll-8 form strong Myo-II cables around them similar to those present at tissue compartment boundaries, while quantitative differences of Toll-8 expression at low levels lead to planar polarity of Myo-II across several cells. Thus, mechanical boundaries and planar polarized mechanical interfaces might be considered as a continuum, depending on quantitative inputs of cell surface protein asymmetries, such as Toll-8, interpreted by adhesion GPCRs, such as Cirl. The role of GPCRs and G protein signaling in conveying planar polarized input quantitatively is substantiated by the observation that overexpression of heterotrimeric G protein subunits Gβ/Gγ increases Myo-II planar polarization in early *Drosophila* embryos and converts intercalating cell columns into boundary like structures (Garcia De Las Bayonas et al., 2019).

Recently Toll receptors, including Toll-8, were shown to be involved in cell competition (Alpar et al., 2018; Meyer et al., 2014) and growth (Germani et al., 2018). Toll-8 is differentially expressed between Myc cells and wild-type cells (Alpar et al., 2018). The interplay between cell competition or tumor growth and mechanics was recently documented (Levayer et al., 2015, 2016; Moreno et al., 2019; Vishwakarma and Piddini, 2020; Wagstaff et al., 2016). It is tantalizing to suggest a role for Cirl and Toll-8 in this context.

The vertebrate Cirl homologues Latrophilins regulate synaptogenesis and neuronal migration via *trans*-heterophilic binding with FLRTs (Boucard et al., 2014; O’Sullivan et al., 2012; Sando et al., 2019; del Toro et al., 2020), which share sequence similarities with Toll-8 (Dolan et al., 2007). FLRT3 plays an important role in tissue morphogenesis (Karaulanov et al., 2006; Smith and Tickle, 2006; Tomás et al., 2011), which might depend on its interaction with Latrophilins.

The depletion of Cirl observed around Toll-8 overexpressing cells resembles the effect of Flamingo (CELSR in vertebrates) overexpression on Frizzled localization (Chen et al., 2008; Strutt and Strutt, 2008), both central components of the conserved core Planar Cell Polarity (PCP) pathway (Devenport, 2014). Intriguingly, Flamingo is also an adhesion GPCR and shares sequence homology with Cirl (Langenhan et al., 2013). It is also required for junctional Myo-II activation in the chick neural plate (Nishimura et al., 2012) and during axis extension in Xenopus (Shindo and Wallingford, 2014). The self-organized planar polarization of Toll-8/Cirl identified in our study suggests the possibility that aGPCRs, such as Cirl and Flamingo, are conserved cell surface proteins involved in PCP that evolved different modalities of symmetry breaking. In the core PCP pathway, Flamingo symmetry breaking is thought to be biased by long range mechanical (Aigouy et al., 2010; Aw et al., 2016; Chien et al., 2015) or chemical gradients of adhesion molecules (Fat, Dachsous) or ligands (Wnt) (Aw and Devenport, 2017), which align Flamingo polarity across the tissue. In the system described here, Cirl symmetry breaking is potentially triggered by quantitative differences in Toll-8 transcriptional levels between neighboring cells. Directionality of spatial differences in Toll-8 transcriptional levels could define the orientation of Toll-8 and Cirl polarity, resembling the Fat-Ds PCP pathway (Bosveld et al., 2012). Thus, our work illustrates a novel protein complex that generates planar polarized mechanical interfaces instructed by tissue-level cues.

**Table 2:** List of genotypes employed in the experiments in the indicated figure panels.

## Author contributions

J.L., Q.M. and T.L. conceived the project. Q.M., J.L. and S.K. performed experiments in embryos, Q.M analyzed them. J.L. performed all the experiments and data analysis in fixed wing discs. S.H. performed and analyzed live experiments in wing discs, and quantified data in Figure 5 and Figure 6. Q.M. and A.L. prepared the samples for the mass spectrometry which was done and analyzed by S.A. and L.C. J-M.P. designed and generated the molecular constructs. J.L., Q.M. and T.L. wrote the manuscript and all authors made comments.

## Supporting information

Table 1_Toll-8_APMS

Table 2_Flies genotype

## Acknowledgments

We thank all members of the Lecuit team, B. Aigouy (IBDM, France) for stimulating discussions during the course of this project, the IBDM imaging facility for microscopy assistance, FlyBase for maintaining curated databases and the Bloomington stock center for providing fly stocks. We thank T. Langenhan and N. Scholz (Leipzig, Germany) for sharing information about Cirl/Latrophilin and fly reagents, and for stimulating discussions. We thank B. Aigouy (IBDM, France) and P. Villoutreix (Centuri, France) for developing the method for quantifying clone smoothness in wing discs. We thank B. Habermann (IBDM, France) for her valuable guidance on performing pair-wise alignment between Toll-8 and FLRTs. We are grateful to M. Ludwig (Birmingham, UK), P.F. Lenne (IBDM, France), E. Wieschaus (Princeton, USA), A. Martin (MIT, USA), R. Karess (IJM, France), G. Jiménez (IBMB, Spain) and T. Gregor (Pasteur Institute, France) for the gift of flies and vectors.

This work was supported by grants from the ERC (Biomecamorph no. 323027) and the Ligue Nationale Contre le Cancer (Equipe Labellisée 2018). J.L. was supported by the Fondation Bettencourt Schueller and the Collège de France. S.H. was supported by an EMBO Long-Term Fellowship (EMBO ALTF 217-2017) and by a Centuri Postdoctoral Fellowship (Centuri, France). The Marseille Proteomics (IBiSA) is supported by Institut Paoli-Calmettes (IPC) and Canceropôle PACA. Proteomics analysis was supported by the Institut Paoli-Calmettes and the Centre de Recherche en Cancérologie de Marseille. Proteomic analyses were done using the mass spectrometry facility of Marseille Proteomics (marseille-proteomique.univ-amu.fr) supported by IBISA (Infrastructures Biologie Santé et Agronomie), Plateforme Technologique Aix-Marseille, the Cancéropôle PACA, the Provence-Alpes-Côte d’Azur Région, the Institut Paoli-Calmettes and the Centre de Recherche en Cancérologie de Marseille. We acknowledge the France-BioImaging infrastructure supported by the French National Research Agency (ANR–10–INBS-04-01, Investments for the future).

## Competing financial interests

The authors declare no competing financial interests.

## Data and materials availability

The authors declare that the data supporting the findings of this study are available within the paper and its supplementary information files. Raw image data are available upon reasonable request. Codes and material are available upon request. The mass spectrometry proteomics data have been deposited to the ProteomeXchange Consortium via the PRIDE partner repository with the dataset identifier PXD017895.

## Movie Legends

**Movie S1:** Time lapse movie of the embryonic ventrolateral ectoderm during axis extension in a control (left) and a *cirl* null mutant (right) embryo, showing cell outlines labeled with LifeAct–mCherry (grey) and tracked cell groups as pseudo-colour overlays. Frames are re-aligned to keep the centre of tracked cell group constant. Framer rate: 30 seconds/frame.

Duration: 20 minutes. Scale bar: 5 μm.

**Movie S2:** Time lapse movie of a wing disc showing live dynamics of Toll-8–YFP (green) and MRLC–Ch (magenta) in a nascent Toll-8–YFP overexpressing clone. Toll-8–YFP starts being detectable at 0 min and its levels increase during the course of the time lapse. Asterisks indicate a cell that dynamically upregulates expression of Toll-8–YFP during ∼50min. Myo-II is being enriched at junctions that show Toll-8–YFP planar polarity (arrow) and this enrichment is lost once Toll-8–YFP expression reaches similar levels between contacting cells (arrowheads). Scale bar: 5 μm.

**Movie S3:** Time lapse movie of a wing disc showing live dynamics of Toll-8–YFP (green) and MRLC–Ch (magenta) in a nascent Toll-8–YFP overexpressing clone. Scale bar: 5 μm. **Movie S4:** Time lapse movie of a wing disc showing live dynamics of Toll-8–YFP (green) and MRLC–Ch (magenta) in Toll-8–YFP overexpressing clones. Scale bar: 5 μm.

---

**Figure S1.**
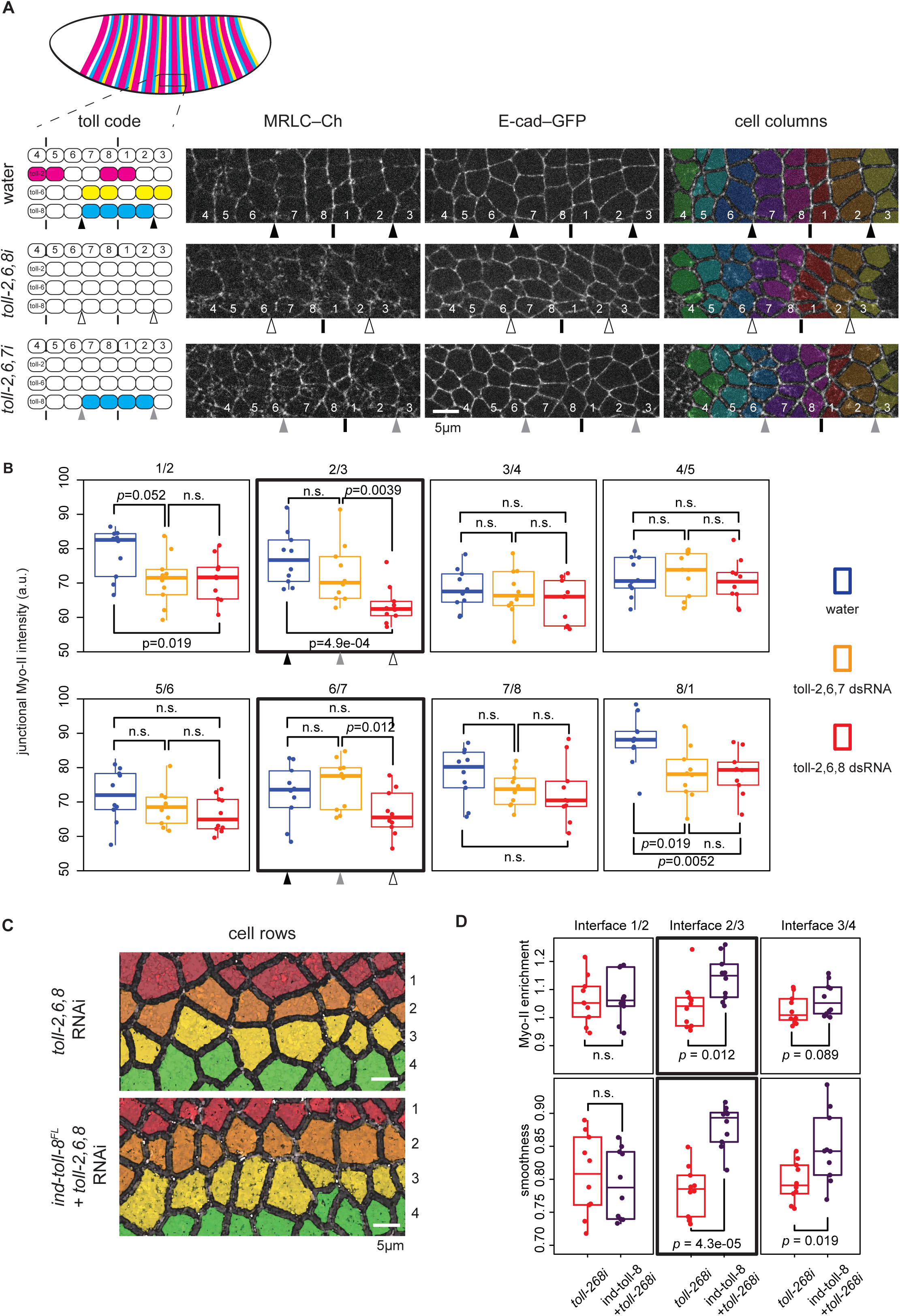
Endogenous Toll-8 is able to enrich Myo-II at its expression boundaries independent of Toll-2,6 (related to Figure 1). **(A)** Schematics of endogenous Toll-2,6,8 expression in embryos under various conditions (top left). Cell columns 1-4: odd parasegments (*even-skipped*+); 5-8: even parasegment (*even-skipped*-). Arrowheads show the two boundaries of Toll-8 expression at positions 2/3 and 6/7. See methods for details. Still images from time lapse movies of embryos injected with water (top), dsRNAs against Toll-2,6,8 (middle) and dsRNAs against Toll-2,6,7 (bottom). Pseudo colors represent cell columns (right). In all panels, vertical bars represent parasegment boundaries where Myo-II is enriched despite of Toll-2,6,8 knockdowns. Arrowheads demark boundaries of the Toll-8 expression domain in embryos injected with water (black), dsRNAs against Toll-2,6,8 (white) or Toll-2,6,7 (grey), where Myo-II is enriched in the sole presence of Toll-8 (bottom). **(B)** Quantifications of junctional Myo-II levels at indicated interfaces in embryos injected with water (blue, *n*=10), dsRNAs against Toll-2,6,7 (orange, *n*=10) and dsRNAs against Toll-2,6,8 (red, *n*=10). Black, white and grey arrowheads show quantifications of representative interfaces highlighted in (**A**). **(C and D)** Additional quantifications for Figures 1F-H. Pseudo colors denote 4 horizontal cell rows in *wt* embryos (top, *n*=10) or embryos expressing ind-Toll-8^FL^–HA (bottom, *n*-10), both injected with dsRNAs against Toll-2,6,8 (**C**). Quantifications of junctional Myo-II enrichment (top) and boundary smoothness (bottom) at interfaces 1/2 (left, interfaces within the ind-Toll-8^FL^– HA domain), 2/3 (center, boundary interfaces) and 3/4 (right, interfaces within the wild-type tissue) (**D**). Scale bars: 5 µm. Statistics: Mann-Whitney U test; n.s.: *p* > 0.05.

**Figure S2.**
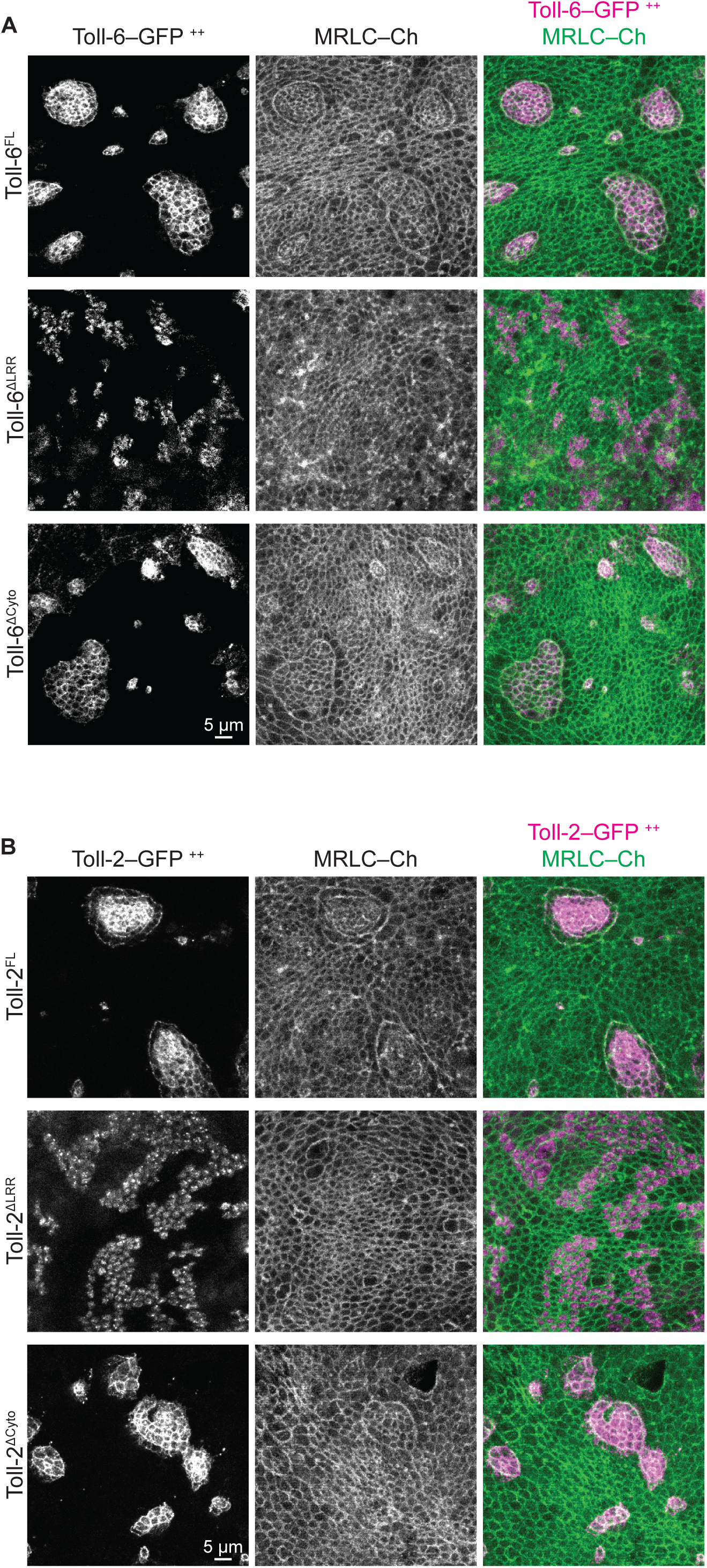
Asymmetric expression of Toll-6 and Toll-2 without their cytoplasmic tails leads to Myo-II enrichment in wing discs (related to Figure 2). **(A)** Fixed Toll-6–GFP (left) and MRLC–Ch (middle) signals from wing disc clones overexpressing full length Toll-6 (Toll-6^FL^, top), Toll-6 with the extracellular LRRs removed (Toll-6^ΔLRR^, middle), or Toll-6 with the intracellular cytoplasmic tail removed (Toll-6^ΔCyto^, bottom). Myo-II enrichment requires the extracellular LRRs but not the cytoplasmic tail of Toll-6. **(B)** Fixed Toll-2–GFP (left) and MRLC–Ch (middle) signals from wing disc clones overexpressing full length Toll-2 (Toll-2^FL^, top), Toll-2 with the extracellular LRRs removed (Toll-2^ΔLRR^, middle), or Toll-2 with the intracellular cytoplasmic tail removed (Toll-2^ΔCyto^, bottom). Myo-II enrichment requires the extracellular LRRs but not the cytoplasmic tail of Toll-2. Scale bars: 5 µm.

**Figure S3.**
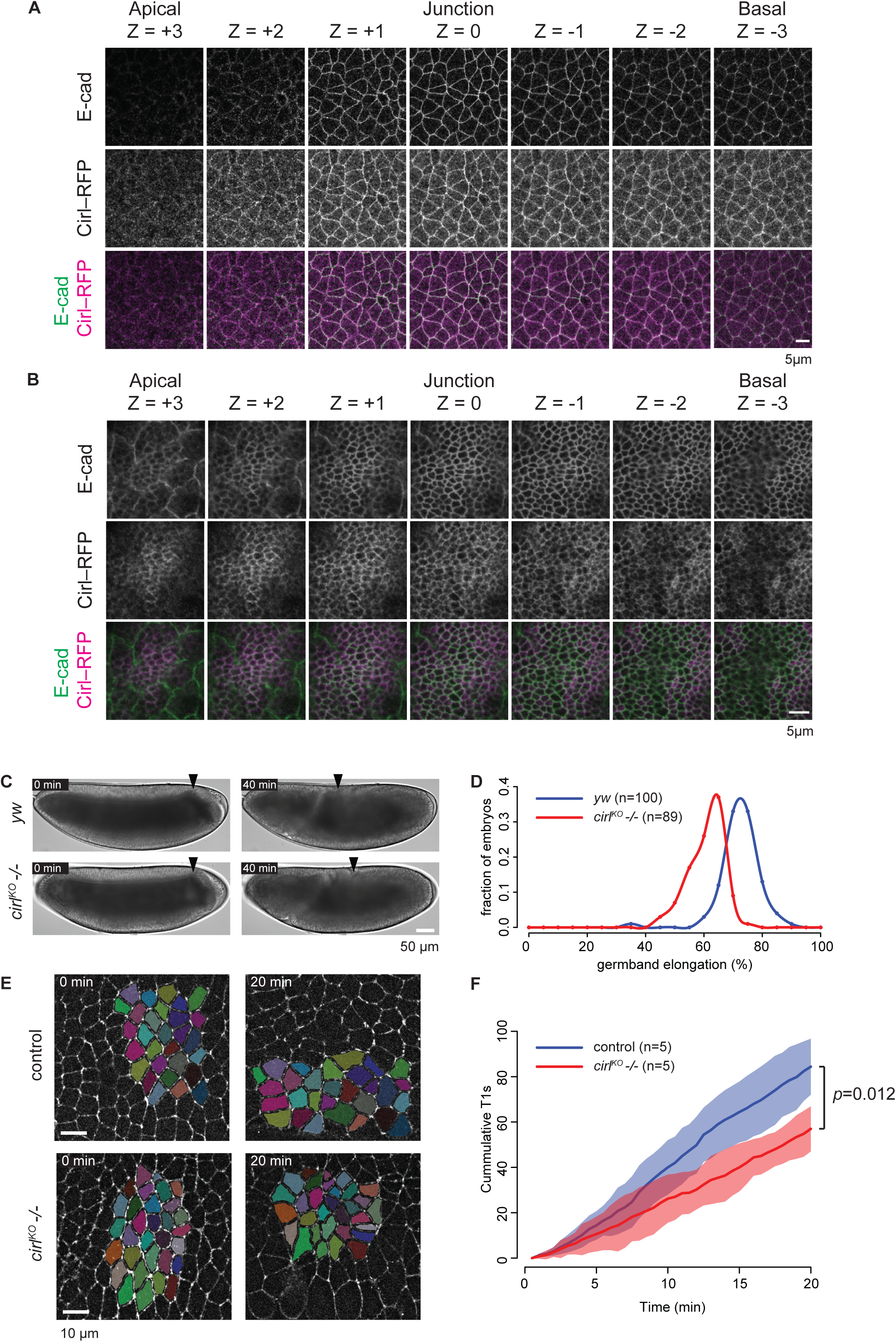
Localization of Cirl in the *Drosophila* embryonic germband and wing disc cells, and its requirement in embryonic axis extension and cell intercalations (related to Figure 3). **(A)** Anti E-cad (green) and anti Cirl–RFP (magenta) signals in the ectoderm epithelium in stage 7 *Drosophila* embryos. Sequential Z planes are shown from apical (left) to basal (right). Step size: 0.38 µm. **(B)** Anti E-cad (green) and anti Cirl–RFP (magenta) signals in the wing disc pouch epithelium. Sequential Z planes are shown from apical (left) to basal (right). Step size: 0.25 µm. In both (**A**) and (**B**), Cirl–RFP is localized at cell-cell interfaces around adherens junctions. **(C)** Axial extension at 0 and 40 minutes in *yw* (top, n=100) and *cirl^KO^ -/-* (bottom, n=89) embryos. 0 min is defined as the beginning of axial extension. Arrowheads denote the dorsal edge of the posterior midgut primordium. Axial extension is slowed down in the absence of *cirl*. **(D)** Histogram of axial extension for *yw* (blue, n=100) and *cirl^KO^ -/-* (red, n=89) embryos for the conditions shown in (**C**). **(E)** Still images from time lapse movies in *wt* (top, *n*=5) or *cirl^KO^ -/-* (bottom, *n*=5) embryos. LifeAct–Ch marks cell outlines. Pseudo colors mark tracked cells. 0 min is defined as the beginning of axial extension. **(F)** Cumulative numbers of T1 transitions in *wt* (blue, *n*=5) or *cirl^KO^ -/-* (red, *n*=5) embryos for conditions shown in (**E**). Solid lines represent mean values, shaded areas represent standard deviation. *p*-value is calculated for total numbers of T1 transitions at the end of 20 minutes between *wt* and *cirl^KO^ -/-* embryos. Scale bars: 5 μm in (**A**) and (**B**); 50 μm in (**C**); 10 μm in (**E**). Statistics: Mann-Whitney U test.

**Figure S4.**
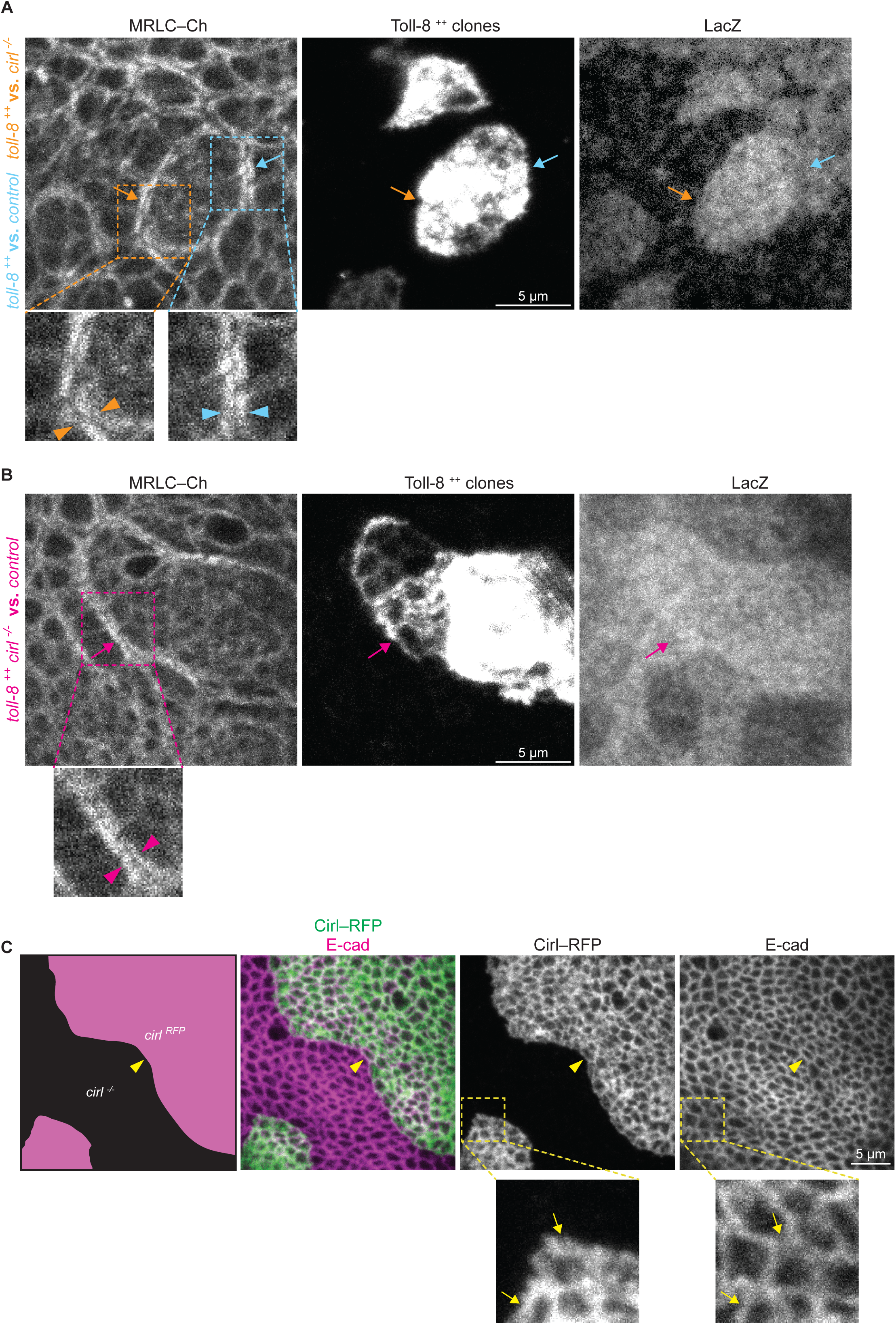
Myo-II is enriched on both sides of the Toll-8 expression boundary (related to Figure 4). **(A)** Myo-II signals in MARCM clones in the wing disc where cells overexpressing Toll-8– YFP (*toll-8 ^++^, lacZ ^+/+^*, green in Figure 4A) are juxtaposed with control cells heterozygous (*cirl ^+/-^, lacZ ^+/-^*, blue in Figure 4A, cyan arrow) or null mutant (*cirl ^-/-^, lacZ ^-/-^*, black in Figure 4A, orange arrow) for *cirl*. The cyan and orange arrowheads on the zoomed images indicate Myo-II enrichment on both sides of clone boundaries (related to Figure 4A). **(B)** MARCM clones in the wing disc. Myo-II is enriched at clone boundaries (magenta arrow) where cells overexpressing Toll-8–YFP and null mutant for *cirl* (*toll-8 ^++^, cirl ^-/-^, lacZ ^+/+^*, green in Figure 4B) are juxtaposed with cells heterozygous (*cirl ^+/-^, lacZ ^+/-^* in blue in Figure 4B) or *wild-type* (*cirl ^+/+^, lacZ ^-/-^* in black in Figure 4B) for *cirl*. The magenta arrowheads on the zoomed image mark Myo-II enrichment on both sides of the clone boundary (related to Figure 4B). **(C)** Cirl–RFP is still localized in wild-type cell interfaces in contact with *cirl* mutant cells (arrowheads and arrows). Scale bars: 5 µm.

**Figure S5.**
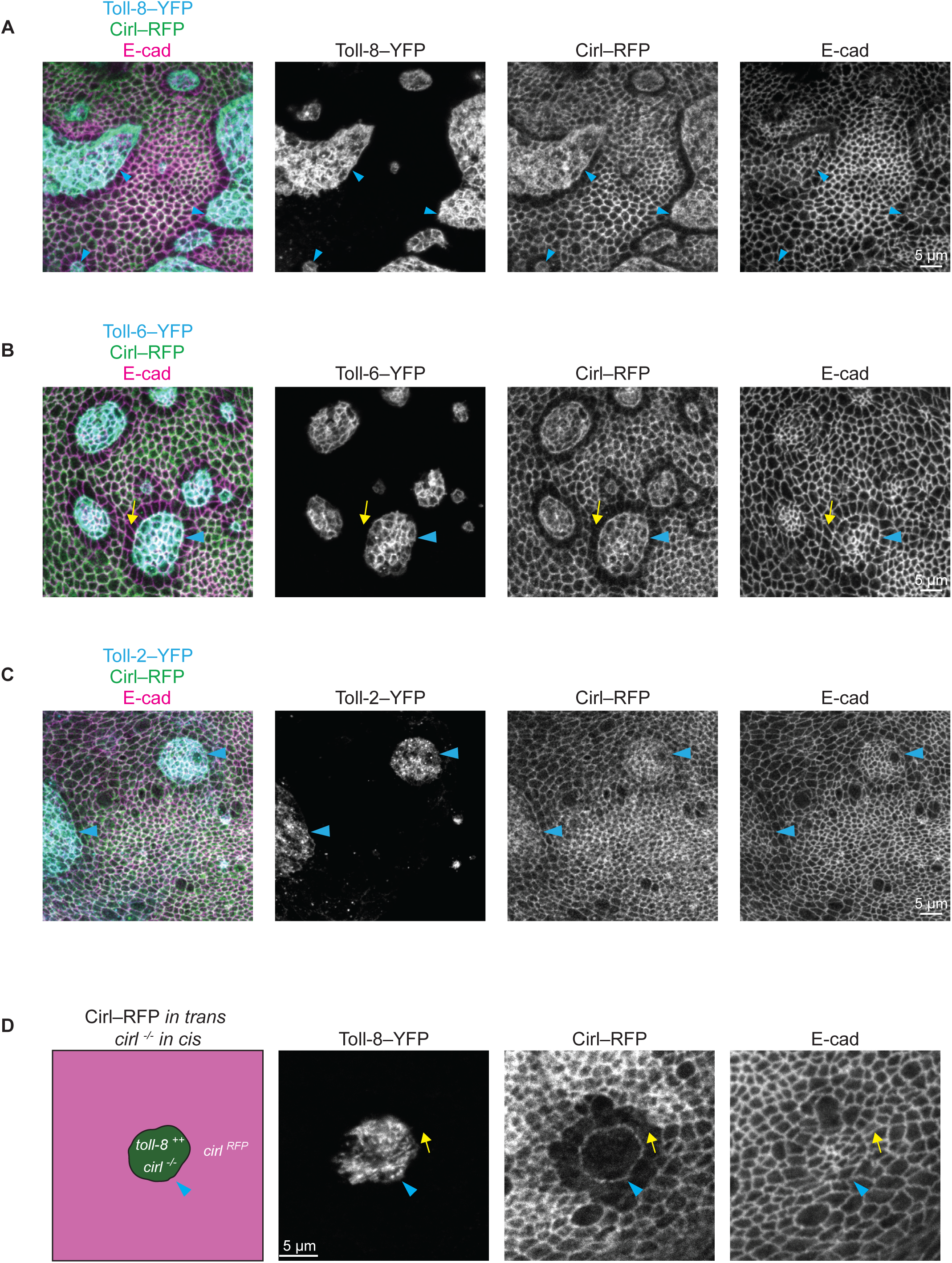
Effect of Toll-8, Toll-6 and Toll-2 overexpression on Cirl distribution in the wing disc (related to Figure 5). Arrowheads indicate clone boundaries in all panels. **(A)** Toll-8–YFP overexpressing clones in a Cirl–RFP wing disc (similar to Figure 5A). **(B)** Toll-6–YFP overexpressing clone in a Cirl– RFP wing disc. Cirl–RFP is depleted from junctions orthogonal to the clone boundary (arrow). **(C)** Toll-2–YFP overexpressing clone in a Cirl–RFP wing disc. Cirl–RFP localization is not affected by Toll-2 overexpression. **(D)** Same experimental setup as in Figure 5B, with the exception that *cirl* is null mutant instead of untagged in Toll-8 overexpressing clones (*toll-8 ^++^*, *cirl ^-/-^* in green). Planar polarity of Cirl in wild-type cells neighboring Toll-8 overexpressing clones is not disturbed by the absence of Cirl inside the clones. Scale bars: 5 μm.

**Figure S6.**
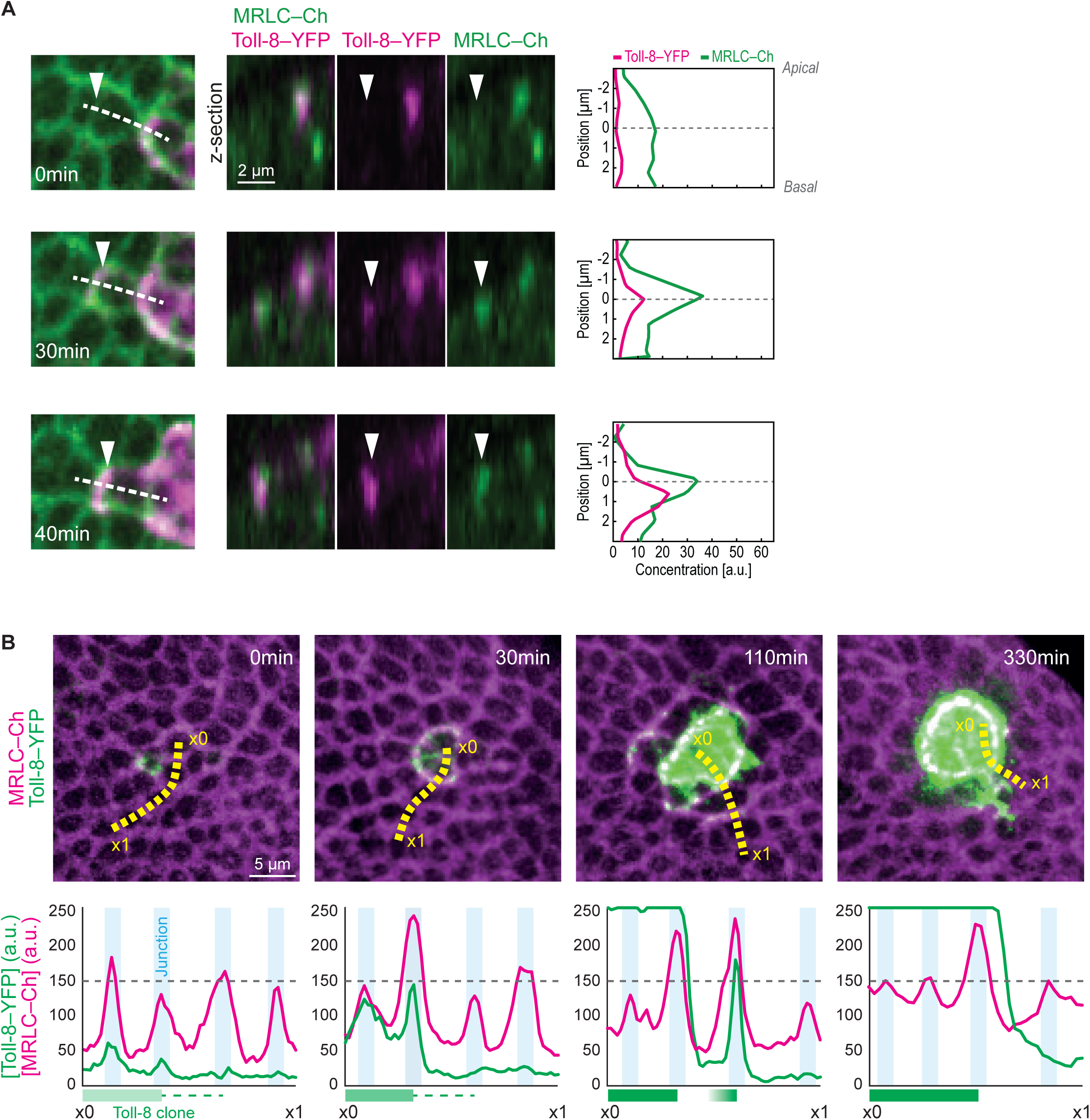
Additional quantifications of nascent Toll-8 overexpressing clones in wing disc movies (related to Figure 6). **(A)** Stills from a movie showing a nascent Toll-8–YFP overexpressing clone in a wing disc (left) and corresponding optical cross-sections (middle) at positions indicated by the dashed lines in the left panels. Quantifications (right) of Toll-8–YFP and MRLC–Ch levels along the apicobasal axis of a cell interface (arrowhead) at subsequent time-points in a cell starting to express Toll-8–YFP at t=∼0min. Myo-II enrichment is observed when Toll-8–YFP is localized exclusively to cell-cell interfaces (t=30min and 40min). **(B)** Quantifications of temporal dynamics of Toll-8–YFP (green) and MRLC–Ch (magenta) at the boundary of a nascent Toll-8–YFP overexpressing clone in the wing disc (panels taken from Figure 6A). Toll-8–YFP and MRLC–Ch levels were quantified along a line (yellow, from x0 to x1) at representative stages of clonal Toll-8 upregulation. Myo-II is enriched at the boundary of Toll-8–YFP expressing cells facing Toll-8–YFP negative cells (middle left). Myo-II is also enriched at interfaces between cells expressing different levels of Toll-8–YFP (middle right) leading to multiple rows of Myo-II enrichment. Once Toll-8–YFP levels equalize between cells, Myo-II only remains enriched at junctions that display differences in Toll-8 expression (right). Scale bars: 5 µm in (**A**) and 2 µm in (**B**).

## Methods

### Fly strains and genetics

The following mutant alleles and insertions were used: *ind-Toll-8^FL^::HA* (attP40 on 2L), *ind- Toll-8^ΔLRR^::HA* (attP40 on 2L), *UASp-Toll-8^FL^::eGFP* (attP40 on 2L), *UASp-Toll-8^ΔLRR^::eGFP* (attP40 on 2L), *UASp-Toll-8^ΔCyto^::eGFP* (attP40 on 2L), *UASt-Toll-8::sYFP2* (VK27 (attP9744) on 3R), *UASp-Toll-6^FL^::eGFP* (attP40 on 2L), *UASp-Toll-6^ΔLRR^::eGFP* (attP40 on 2L), *UASp-Toll-6^ΔCyto^::eGFP* (attP40 on 2L), *UASt-Toll-6::sYFP2* (VK27 on 3R), *UASp-Toll-2^FL^::eGFP* (attP40 on 2L), *UASp-Toll-2^ΔLRR^::eGFP* (attP40 on 2L), *UASp-Toll-2^ΔCyto^::eGFP* (attP40 on 2L), *UASt-Toll-2::sYFP2* (attP40 on 2L), *Cirl^KO^* (LAT84 and LAT154, gifts from T. Langenhan) (Scholz et al., 2015), *Cirl::RFP* knock-in (Scholz et al., 2017) (*Cirl::RFP^KIN^*, LAT159, gift from T. Langenhan), *sqh-Lifeact::mCherry* (VK27 on 3R, gift from P.F. Lenne), *E-cad::eGFP* knock-in (Huang et al., 2009) (*E-cad::eGFP^KIN^*), *eve::sYFP2^BAC^* (BAC construct, S2E.MSE.eve.YFP, FBal0279504, gift from M. Ludwig) (Ludwig et al., 2011), *hs-FLP* (Chan et al., 2017), *Ubx-FLP* (Bloomington BL 42718), *Act5C>STOP>GAL4* (Bloomington BL 4780), *FRT42D* (Bloomington BL 1802), *FRT42D arm-LacZ* (Bloomington BL 7372), *FRT42D tub-GAL80* (Bloomington BL 9917), *tub-GAL80ts* (Bloomington BL 7108) and *tub-GAL80ts* (Bloomington BL 7017). *67-Gal4* ({matαtub-GAL4-VP16}67) is a ubiquitous, maternally supplied GAL4 driver (gift from E. Wieschaus). MRLC is encoded by the gene *spaghetti squash* (*sqh*, Genebank ID: AY122159). Sqh was visualized using *sqh-Sqh::mCherry* (VK18 (attP9736) on 2R or VK27 (attP9744) on 3R for experiments in the wing disc or a construct on chromosome 2 from A. Martin (Martin et al., 2009) for live-imaging experiments in the embryo) and *sqh-Sqh::eGFP* transgenics (gift from R. Karess).

All fly constructs and genetics are listed in Table 2.

### Constructs and transgenesis

#### Expression vectors drivers

UASp expression vector driver was generated by inserting a PhiC31 attB sequence downstream from the K10 3’-UTR of pUASp. UASt expression vector driver corresponds to pUASTattB which contains a PhiC31 attB sequence inserted downstream from the SV40 3’-UTR (Bischof et al., 2007).

*ind* is an early horizontal stripe expression vector driver generated by modifying the *pbphi-evePr-MS2-yellow* (Bothma et al., 2014) (gift from T. Gregor) as follows. First, EVEstr2 enhancer sequence was replaced by the ind^1.4^ enhancer (Lim et al., 2013) (gift from Gerardo Jiménez). Second, a part of Hsp70Bb 5’-UTR was added after the eve basal promoter. Third, MS2-yellow sequence was replaced by a small polylinker for further cloning.

#### Expression vectors constructs and transgenics

*Toll-8* (*Tollo*, CG6890), *Toll-6* (CG7250) and *Toll-2* (18 wheeler, CG8896) whole ORFs were amplified using specific pACMAN genomic plasmids and cloned inside each expression vectors. UASp driven Tolls ORFs were all tagged Cterminally by mEGFP with a GSAGSAAGSGEF flexible aa linker in between*.UASp-Toll-8^FL^::eGFP* is the full length Toll-8 (1346aa). *UASp-Toll-8^ΔCyto^::eGFP* is a cytoplasmic truncated version of this vector (deletion from aa H1052 to M1346 last aa). In *UASp-Toll-8^ΔLRR^::eGFP*, all LRR repeats were removed (deletion from aa E99 to L917). *UASp-Toll-6^FL^::eGFP* is the full length Toll-6 (1514aa). *UASp-Toll-6^ΔCyto^::eGFP* is a cytoplasmic truncated version of this vector (deletion from aa H1088 to A1514 last aa). In *UASp-Toll-6^ΔLRR^::eGFP*, all LRR repeats were removed (deletion from aa A139 to G964). *UASp-Toll-2^FL^::eGFP* is the full length Toll-2 (1385aa). *UASp-Toll-2^ΔCyto^::eGFP* is a cytoplasmic truncated version of this vector (deletion from aa F1026 to V1385 last aa). In *UASp-Toll-2^ΔLRR^::eGFP*, all LRR repeats were removed (deletion from aa F110 to L900). *UASt-Toll-2::sYFP2, UASt-Toll-6::sYFP2* and *UASt-Toll-8::sYFP2* are Cter sYFP2 tag construct of full length Toll-2,6 and 8 ORFs cloned into UASt using the same GSAGSAAGSGEF flexible aa linker in between. *ind-Toll-8^FL^::HA* is a Cter HA tag construct of full length Toll-8 with no linker in between. In *ind-Toll-8^ΔLRR^::HA*, all LRR repeats were removed (deletion from aa E99 to L917).

All recombinant expression vectors were built using “In-Fusion cloning” (Takara Bio), verified by sequencing (Genewiz) and sent to BestGene Incorporate for PhiC31 site specific mediated transgenesis into attP40 (2L, 25C7) or VK27 (attP9744, 3R, 89E11). Fully annotated FASTA sequences of all these vectors are available on request.

### Antibodies

The following primary antibodies were used: rat-anti-E-Cad (1:200, DHSB DCAD2 concentrate), mouse-anti-β-catenin (1:400, DHSB N2 7A1 Armadillo concentrate), mouse-anti-LacZ (1:100, DHSB 40-1a concentrate), rat-anti-HA (1:100, Anti-HA High Affinity rat IgG_1_, Roche ROAHAHA). Sqh::eGFP was detected with rabbit-anti-GFP (1:500, Life Technologies A11122 or 1:1000 Abcam ab6556). Cirl::RFP was detected with rabbit-anti-RFP (1:1000, Rockland 600-401-679). The following secondary antibodies were used: donkey-anti-rabbit Alexa Fluor 488 IgG (Life Technologies A 21206), donkey-anti-rabbit Alexa Fluor 568 IgG (Life Technologies A10042), donkey anti-mouse Alexa Fluor 568 IgG (Life Technologies A10037), donkey-anti-mouse Alexa Fluor 647 IgG (Jackson ImmunoResearch 715 605 151) and donkey-anti-rat Alexa Fluor 647 IgG (Jackson ImmunoResearch 712 605 153). All secondary antibodies were used at 1:500.

### Affinity Purification Mass Spectrometry

#### Protein purification and mass spectrometry

Roughly 600 embryos for each sample were collected from overnight cages kept at 25°C for the following crosses: *yw* (control) or females *; 67-GAL4/+; UASt-Toll-8::sYFP2/+;* x males *; 67-GAL4/+; UASt-Toll-8::sYFP2/+;* (Toll-8::YFP maternal and zygotic overexpression), dechorionated with bleach, transferred directly to lysate buffer (10 mM Tris/Cl pH 7.5; 150 mM NaCl; 0.5 mM EDTA; 0.5% NP-40, supplemented with protease and phosphatase inhibitors) and crushed manually on ice over 30 minutes. Lysates were centrifuged to clear debris and protein concentrations of post-centrifugation supernatants were determined. The crude protein yield per lysate sample is usually 1000∼3000 µg. In each experiment, lysates of comparable protein concentration were incubated with pre-rinsed GFP nano-trap agarose resin (Chromotek, gta-20) at 4°C for 90 min, rinsed 3 x and resuspended in 2x Laemmli buffer with DTT. Protein extraction and purification was performed 3 times each for each cross and verified with silver staining. Protein samples were further purified on NuPAGE 4-12% Bis-Tris acrylamide gels (Life Technologies) and treated with in-gel trypsin digestion (Shevchenko et al., 1996) with minor modifications. Peptides were harvested with two extractions, first in 5% formic acid and then in 5% formic acid in 60% acetonitrile. Samples were reconstituted with 0.1% trifluoroacetic acid in 4% acetonitrile and analyzed by liquid chromatography (LC)-tandem mass spectrometry (MS/MS) with an Orbitrap Fusion Lumos Tribrid Mass Spectrometer (Thermo Electron, Bremen, Germany) online with an Ultimate 3000RSLCnano chromatography system (Thermo Fisher Scientific, Sunnyvale, CA). A detailed mass spectrometry protocol is available upon request.

#### Protein identification and quantification

Relative intensity-based label-free quantification (LFQ) was processed using the MaxLFQ algorithm (Cox et al., 2014) from the freely available MaxQuant computational proteomics platform (Cox and Mann, 2008). Spectra were searched against a *Drosophila melanogaster* database (UniProt Proteome reference, date 2017.08; 21982 entries). The false discovery rate (FDR) at the peptide and protein levels were set to 1% and determined by searching a reverse database. For protein grouping, all proteins that could not be distinguished based on their identified peptides were assembled into a single entry according to the MaxQuant rules. The statistical analysis was done with Perseus program (version 1.5.1.6) from the MaxQuant environment (www.maxquant.org). Quantifiable proteins were defined as those detected in at least 67% of samples in at least one condition. Protein LFQ normalized intensities were base 2 logarithmized to obtain a normal distribution. Missing values were replaced using data imputation by randomly selecting from a normal distribution centered on the lower edge of the intensity values that simulates signals of low abundant proteins using default parameters (a downshift of 1.8 standard deviation and a width of 0.3 of the original distribution). To determine whether a given detected protein was specifically differential, a two-sample t-test was done using permutation-based false discovery rate (pFDR) with a threshold at 0.1% (5000 permutations). The *p*-value was adjusted using a scaling factor s0=1 (Table 1). In Figure 3, differential proteins are highlighted by a cut-off for log2|Fold change|>2 and a *p*-value<0.01. The mass spectrometry proteomics data have been deposited to the ProteomeXchange Consortium via the PRIDE (Vizcaíno et al., 2014) partner repository with the dataset identifier PXD017895.

### Bright-field live imaging in embryos

Images of wild-type or mutant embryos were collected on an inverted microscope (Zeiss, AxioVision software) equipped with a programmable motorized stage to record different positions over time (Mark&Find module from Zeiss). Images were acquired every 2 min for 60 minutes from post dorsal movement of the posterior midgut primordium (0 min). The extent of elongation was measured by dividing the travel distance of the posterior midgut primordium at 40 min and normalized to the maximum travel distance.

### RNA interference in embryos

#### dsRNA probes

dsRNA probes were made using PCR product containing the sequence of the T7 promoter (TAATACGACTCACTATAGG) followed by 18-21 nucleotides specific to the gene. The dsRNA probe against Toll-2 (18w, CG8896) is 393-bp long and located in the 5’UTR region (Forward primer: AGTTTGAATCGAAACGCGAGGC; Reverse primer: ATGCCAGCCACATCTTCCA). The dsRNA probe against Toll-6 (CG7250) is 518-bp long and located in the 5’UTR region (Forward primer: TCGAAAATCAGCCAACGTGC; Reverse primer: CGATTCACGGTTTAGCTGCG). The dsRNA probe against Toll-7 (CG8595) is 749-bp long and located in the coding region (Forward primer: TGGCAACCGTCTGGTTACTC; Reverse primer: CGTTCATGATGCTCTGCGTG). The dsRNA probe against Toll-8 (Tollo, CG6890) is 423-bp long and located in the 5’UTR region (Forward primer: CGTTTGTCGTTCAGCGGATG; Reverse primer: CCCCTCATAACCTCCCCGAT) and does not target the *ind-Toll-8::HA* transgenes. Gel purified PCR products were subsequently used as a template for the *in vitro* RNA synthesis with T7 polymerase using Ribomax (Promega, P1300). The dsRNA probes were purified using Sure-Clean (Bioline, BIO-37047). Triple dsRNA probes against Toll-2,6,8 and Toll-2,6,7 were prepared and injected at a final concentration of 5 µM each in RNAse-free water.

#### Embryo injections

Embryos were collected from fresh agar plates in cages kept at 25°C allowed for 30-min egg laying. Embryos were then dechorionated in 50% bleach, rinsed and aligned on cover slips (#1.5) covered with heptane-glue. After a few minutes of desiccation, embryos were covered with Halocarbon 200 oil and injected with dsRNA or RNase-free water. Post-injection embryos were stored at 25°C until live imaging.

### Fluorescence live imaging and image processing in embryos

Embryos were aligned on cover slips (#1.5) with heptane-glue and were covered with Halocarbon 200 oil. Dual channel time-lapse imaging was performed on a Nikon Eclipse Ti inverted spinning disc microscope (Roper) with a 100x/1.4 oil-immersion objective at 22°C, controlled by the Metamorph software. Z stacks (step size: 0.5 µm) of 6∼10 slices were acquired every 30 seconds, for 15∼45 minutes starting from stage 6. Laser power was measured and kept constant across all experiments.

To generate 2D projections in experiments with E-Cad::GFP (Figures 1F, S1A and S1C), a custom FIJI macro (Bailles et al., 2019) integrating the ‘stack focuser’ plugin from M. Umorin was used to perform maximum intensity projection for all channels with 3 Z planes around the junctional plane (labeled by E-cad::GFP). For Figure 3C, a single plane at the junction level is manually selected based on maximum junctional sqh::GFP signals. The resulting 2D images were subjected to a background subtraction procedure using the rolling ball tool (radius 50 pixels). The 2D images were segmented on E-cad::GFP or LifeAct::Ch channels semi-automatically with manual corrections in the FIJI plug-in Tissue Analyzer (Aigouy et al., 2010). The resulting segmentation masks were then dilated by 5 pixels on either side of the junction and used as masks for subsequent quantifications. Cell tracking and quantifications of T1 transitions are performed semi-automatically with Tissue Analyzer with manual correction.

### Immunofluorescence and image processing in embryos

Embryos were fixed with 8% formaldehyde for 20 min at room temperature. Embryos were processed and stained according to standard procedures (Müller, 2008). Embryos were mounted in Aqua-Polymount (Polysciences). Images were acquired on a Leica SP8 inverted confocal microscope with a 63x/1.4 NA oil-immersion objective (with exception of Figure S3A acquired on a Zeiss LSM780 with a 63x/1.4 NA oil-immersion objective). Z stacks with step size of 0.25-0.4 μm were collected.

2D images were generated by maximum intensity projections followed by the same procedure as for live imaging experiments in embryos (except for 3-pixel dilatation in segmentation masks generated from β-catenin stainings).

### Clonal analysis in wing discs

Flies were allowed to lay eggs in vials for ∼8h at 25°C and vials were kept at 25°C until heat-shock.

For clonal overexpression of Tolls (Figures 2, 3I, 5A, S2 and S5A-S5C) 72h AEL (after egg laying) old larvae were heat-shocked at 37° for 10-14 minutes and dissected after 24h. For GAL80ts experiments (Figures 6 and S6), 72h AEL larvae were heat-shocked at 37° for 12 minutes, kept at 18° for 48 hours and subsequently incubated at 30°C for 2h15min in order to inactivate GAL80ts and allow expression of Toll-8::YFP.

For MARCM experiments (Mosaic Analysis with a Repressible Cell Marker, Figures 4A, 4B, 5B, 5C, S4A, S4B and S5D), 72h AEL larvae were heat-shocked at 37° for 1h, kept at 18° and heat-shocked again 7 hours later at 37° for 1h. Larvae were kept at 18° for 20 hours, shifted to 25° and dissected 24h later. Keeping the larvae at 18° allowed growth of the clones in the presence of no/low levels of Toll-8 expression. Larvae to observe *cirl* mutant clones (Figure 4D) were treated the same way.

### Immunofluorescence and image processing in wing discs

Staged larvae were dissected in PBS, transferred to 4% PFA in PBS and fixed under agitation for 18 min at room temperature. After fixation, wing discs were first rinsed with PBS, then extensively washed with PBT (PBS plus 0.2% Triton-X100) and blocked in PBT with 5% normal donkey serum (NDS, Jackson Immuno Research Laboratories, 017-000-001) for at least 30 min at room temperature, followed by incubation with primary antibody in 2% NDS overnight at 4 °C. The next day wing discs were washed in PBT and incubated in secondary antibody with 2% NDS for 1h30min at room temperature. After six rounds of washes with PBT, samples were mounted in Mowiol (Sigma-Aldrich, 324590). Larval mouth hooks were used as spacers in the experiments where Myo-II was observed. Images were acquired on a Leica SP8 inverted confocal microscope with a 63x/1.4 NA oil-immersion objective. Toll-8::YFP and Sqh::Ch were visualized with their endogenous fluorescence. Image stacks with step size of 0.25-0.5 μm were collected.

Peripodial signal was masked from the image stacks in ImageJ to avoid interference with signals from the wing disc proper. 2D projections were generated using the aforementioned custom stack focuser macro in ImageJ, projecting two z planes around the junctional plane of each cell (detected by E-cad staining, except Figures 4A and 4B, projected on Sqh::Ch signals). This allows to project the entire wing pouch independently of the shape of the wing disc. The 2D-projected stacks were then segmented on E-cad stainings (except Figures 4A and 4B, segmented on Sqh::Ch signals) using Tissue Analyzer (Aigouy et al., 2010).

### *Ex vivo* live imaging and image processing in wing discs

The culture medium used for long-term time lapse imaging of wing imaginal disc explants is described in Dye *et al*. (Dye et al., 2017). In short, Grace’s insect medium (Sigma G9771, without sodium bicarbonate) was buffered with 5mM BisTris and the pH adjusted to 6.6-6.7. Subsequently the medium was sterile filtered (0.2µm pore size) and kept at 4°C for up to 4 weeks. At the day of the experiment the medium was supplemented with 5% fetal bovine serum (FBS), Penicillin-Streptomycin (final 1x from a 100x stock solution, Sigma P4333) and warmed to 30°C in a water bath. Just before dissection of the larvae, 20-Hydroxyecdysone (20E, Sigma, H5142) was added to yield a total concentration of 20nM. 20E was kept as a 1000x stock solution in ethanol at -20°C. For the experiment, 72h AEL larvae were heat-shocked at 37° for 12 minutes, kept at 18° for 48 hours and subsequently incubated at 30°C for 2h15min in a water bath. Subsequently, larvae were floated out of the food using 30% glycerol and washed in sterile water twice. Surface sterilization in 70% Ethanol was followed by another wash in sterile water and then in medium. Larvae were dissected in culture medium, wing discs isolated and mounted on a round cover slip. In order to restrict disc movement during imaging, discs were covered by a porous membrane (Whatman cyclopore polycarbonate membranes; Sigma, WHA70602513) using two stripes of double-sided tape as spacers. Finally, this sandwich was mounted in an Attofluor cell chamber (A7816, Invitrogen) and filled with 1ml of medium and covered with Parafilm M (P7793, Sigma-Aldrich) to avoid evaporation. Discs were imaged on a Nikon Eclipse Ti inverted spinning disc microscope (Roper) equipped with an incubation chamber heated to 30°C. Imaging was done using a 60x/1.2 NA water-immersion objective (Figures 6A and S6; Movies S2 and S3) or a 100x/1.4 NA oil-immersion objective (Movie S4). Dual imaging of Toll-8::YFP and Sqh::Ch was performed by simultaneous excitation of fluorophores with 515nm and 561nm laser lines using a dichroic mirror to image on two cameras (Evolve 512, Photometrics). Stacks of 40 slices with 0.7µm spacing were acquired every 10min (60x movies) or every 5min (100x movies).

A maximum projection of the disc proper junctional plane was obtained by masking the peripodial epithelium and the lateral portion of the disc proper manually in ImageJ based on sqh::Ch signals. Background subtraction was done using a rolling ball (50px radius) in ImageJ.

### Data analysis

#### Definition of expression interfaces

When references channels (Toll-8::HA, Toll-8::YFP, Toll-8::GFP, or LacZ staining) were available, expression interfaces were defined from reference channels in Tissue Analyzer.

To define horizontal cell rows in Figures 1F and S1C, cell rows were counted from the ventral midline, with the 4^th^ cell row (most ventral) being 2 rows away from the ventral midline. The boundary between the 2^nd^ and the 3^rd^ cell rows is consistent with the position of *ind* ventral expression boundary.

To define vertical cell columns in Figures S1A and S1B, parasegment boundaries were visualized with Eve::YFP, with the anterior boundary of Eve::YFP signal defined as the parasegment boundary between even- and odd-numbered parasegments. Thus, cell columns 1-4 belong to odd-numbered parasegments (Eve::YFP+), while 5-8 belong to even-numbered parasegments (Eve::YFP-).

#### Quantification of junctional Myo-II intensities

Raw pixel intensities from segmented junctions were measured in Tissue Analyzer. To extract data tables containing raw pixel intensities from Tissue Analyzer, a customized R procedure was developed using the RSQLite package. Adjusted junctional pixel intensities were obtained by subtracting mean cytoplasmic intensity value measured on each image. Enrichment was calculated as ratios of adjusted junctional intensity values between junctions of interest and those in nearby wild-type tissues (Figures 1A and 2A).

#### Quantification of boundary smoothness

Boundary smoothness for ventral *ind* expression boundary in the embryo was calculated as the ratio between distance between two end vertices over total junctional length (Figure 1A). Boundary smoothness value approaches 1 as the boundary gets smoother.

For clone smoothness in the wing disc, an original method developed by P. Villoutreix (Centuri, France) was implemented in Tissue Analyzer by B. Aigouy (IBDM, France) under the plugin ‘Clone wiggliness’. In brief, the boundary of the clone was extracted, the vertices present at the clone boundary were ordered, and an angle was calculated for each vertex with its two neighboring vertices present at the clone boundary (Figure 2A). A mean value per clone was then calculated and this value is getting closer to 180° if the clone is smooth.

#### Quantification of apical-basal protein localization (Figure 5)

Protein localization and concentration along the apical-basal axis was quantified as described in Harmansa *et al*. (Harmansa et al., 2017) from high-z-resolution image stacks (0.35µm slice spacing, acquired on a Leica SP8 confocal microscope using a 63x/1.4 NA oil-immersion objective) of wing discs expressing Toll-8::YFP, Cirl::RFP and stained for the junctional marker E-Cadherin (E-cad). Optical cross-sections were obtained by using the ‘reslice’ option in Fiji software (ImageJ; National Institute of Health) and subsequently background was subtracted (‘rolling ball’ radius of 50px). From these cross sections, junctional fluorescence intensity profiles of E-cad, Toll-8::YFP and Cirl::RFP were extracted along a line of 0.6µm width (corresponding to approximately the width of a junction) using the ‘plot profile’ function in Fiji. Average profiles from different junctions/discs were computed by using the peak of the E-cad profile to align individual profiles. Average profiles were calculated in Excel software (Microsoft) and plotted in Python software using the Seaborn library (line plot function, error bands show the 95% confidence interval computed by bootstrapping). In the plots the junctional plane is visualized by a blue band that is defined as the 0.5µm above and below the average E-cad peak.

#### Quantification of Toll-8 and Cirl planar polarity (Figure 6B-6E)

We restricted our analysis to single cells expressing low Toll-8::YFP levels in the vicinity of a high Toll-8::YFP expressing cell. This ensured that each junction included in our computation only contained Toll-8::YFP originating from the single cell that was quantified. For each single cell, junctional Toll-8::YFP levels were extracted (using the segmented line tool in ImageJ, line width = 6px) along the parallel junctions (Toll-8^‖^, the junctions not being in contact with the high Toll-8 expressing cell) and along the two junctions being in direct contact with high Toll-8 expressing cell (Toll-8^Ʇ^). Toll-8::YFP enrichment at parallel junctions was computed by calculating the ratio between mean Toll-8^‖^ and mean Toll-8^Ʇ^ levels (Figure 6E, scheme). Cirl planar polarity has been computed in an analogous way (Figure 6C, scheme). To test for a correlation between Cirl and Toll-8 planar polarities, polarity values for each doublet of cell was plotted and linear fitting was performed using Python (Seaborn library for plotting and the Stats library to compute the least square regression and *p*-value).

#### Data visualization

With the exception of Figures 5, 6C and S6, data visualization was performed in R with customized scripts. The following custom packages were used: “fields” and “ggplot2”. Plots shown in Figures 5, 6C and S6 were plotted in Python using the Seaborn library.

Box-plot elements are defined as follows: center line represents the median; box limits represent the first and third quartiles (the 25th and 75th percentiles); upper whisker extends from the box limit to the largest value no further than 1.5x interquartile range from the box limit and lower whisker extends from the box limit to the smallest value at most 1.5x interquartile range from the box limit; all data points are plotted on the graph.

### Statistics

*p*-values were calculated using the Mann-Whitney U test in R, except from Figure 5A’’ (two-sided *t*-test with unequal variance) and Figure 6C (two-sided *t*-test performed using the Python Stats library). No statistical method was used to predetermine sample size. The experiments were not randomized, and the investigators were not blinded to allocation during experiments and outcome assessment.

### Repeatability

All measurements were performed in 8–100 independent experiments. Each embryo and clone in the wing disc is considered as an independent experiment.

## Notes

### Competing Interest Statement

The authors have declared no competing interest.

### Summary of Updates

Additional quantifications; Text modifications; Reformating of the figures.

